# Small Extracellular Vesicles from Human Bone Marrow Mesenchymal Stromal Cells enhance migration and regulate gene expression of reparative mechanisms in human dermal fibroblasts

**DOI:** 10.1101/2024.07.19.601214

**Authors:** Rui Lei, Tamiris Borges da Silva, Zhanfeng Cui, Hua Ye

## Abstract

Studies have showed that mesenchymal stromal cells (MSCs) could secrete a variety of bioactive particles, including small extracellular vesicles (sEVs), a type of membrane bound vesicle produced by cells, that might be a key intermediate to the beneficial paracrine effects of MSC therapy. In this study, we harvested the conditioned medium (CM) from both human bone marrow mesenchymal stromal cells (hBM-MSCs) and normal human dermal fibroblasts (NHDFs), fractionated CM of each into the sEV fraction and non-small extracellular vesicle (NsEV) fraction and compared their functions on NHDF-based migration and proliferation assays, as crucial processes in wound healing, coupled with transcriptomic analysis through mRNA sequencing to assess the regulation on gene expression of the recipient NHDFs by secretome exposure. Our findings show that both sEV and NsEV as well as CM from hBM-MSCs showed an overall promotive effect on migration behaviour of NHDFs, but MSC-sEV seems to be more potent for NHDF migration among different MSC secretome fractions and is also superior in effect to the corresponding sEVs produced by the recipient NHDFs (HDF-sEV). Gene ontology analysis revealed enrichment of pathways related to migration, corroborating *in vitro* results; changes in genes within the regulation of proliferation pathway were also significant. Our study provides referential significance for the choice of cellular secretome fractions to be used in wound healing studies and trials. The secretome fractionation and functional comparison strategy using cell-based model and comparative transcriptomics analysis would be a useful tool for comprehensive functional evaluation of cellular secretome.

## Introduction

Mesenchymal stromal cells (MSCs) are self-renewing multipotent cells exhibiting multilineage differentiation and have been used as a powerful therapeutic tool for various human diseases^1^. Bone marrow derived mesenchymal stromal cells (BM-MSCs) have been regarded as a promising source of human MSC-based therapeutic strategy in the field of tissue engineering and regenerative medicine^2,3^. Despite their cell replacement and multipotential differentiation capacity, several studies showed that MSCs have a low engraftment rate in injured area but long-remained therapeutic effects after implantation, suggesting that paracrine factors might be the principal mechanism that contributes to tissue repair^4,5^. Indeed, conditioned medium (CM) harvested from BM-MSCs (MSC-CM) was reported to have promising pharmaceutical potentials in cutaneous regeneration and wound healing in skin models^6–10^, but also BM-MSC secreted small extracellular vesicles (MSC-sEVs), a type of membrane-bound vesicles with a diameter of 30-200 nm, was showed to display regeneration effects on wound healing, promoting proliferation and migration of skin cells, angiogenesis, granulation tissue formation, collagen synthesis and immune regulation^6,11–13^, by mediating cell-cell communication.

It is reasonable to hypothesize that the MSC secretome (MSC-CM) may contain both beneficial factors including soluble proteins and sEVs but also deleterious factors such as metabolism waste that may mediate contradictory effects. However, to our knowledge there are no published efforts that have methodically assessed the functionalities of different fractions of MSC secretome. Therefore, it is necessary to conduct a comprehensive comparison between the functions of MSC-sEV fraction, MSC non-sEV fraction (MSC-NsEV) and the whole MSC-CM itself on tissue repair and wound healing mechanisms through both in vitro cell assay and comparative transcriptomics analysis, with the parallel control of corresponding secretome fractions from recipient cells, to fill the knowledge gap.

Dermal fibroblasts are reported to be stimulated to proliferate and migrate under the stress condition of wound healing, and play a vital role in both inflammatory and proliferative phase, therefore, dermal fibroblast-based skin model has been widely used to test the therapeutic effects of cellular secretome on tissue repair and wound healing process. Shabbir et al. demonstrated a dose-dependent enhancement of proliferation and migration of fibroblasts treated with human bone marrow mesenchymal stromal cell derived sEVs by activating several important signalling pathways (Akt, ERK, and STAT3) in wound healing ^14^. This study was performed with fibroblasts from chronic wound edges and non-defective fibroblasts, showing that MSC secretome was able to promote a reparative effect on both types of fibroblasts. As wound healing is a very complex process, challenging to be faithfully mimicked with 2D culture ^15^, it is not within the scope of this study to assess wound healing in its entirety, but by investigating some of the involved processes (i.e. migration and proliferation).

In this study, we produced and fractionated the conditioned medium of BM-MSCs into sEV fraction and non-sEV fraction, developed in vitro normal human dermal fibroblasts (NHDFs) based proliferation and migration assay to generally assess the functionality of MSC secretome on normal human dermal fibroblasts and enable comparative transcriptomics analysis to reveal hMSC secretome mediated functions on tissue repair, with a proper control of the corresponding secretome fractions from the recipient NHDFs (Figure 1). By investigating the overall impact of different secretome fractions on the cellular response and the transcriptome of the recipient cells, we can identify critically impacted cellular pathways involved in the migration and proliferation processes and obtain an insight into the mechanisms of action of the MSC secretome, which is relevant to uncover EVs mechanisms of action and aid in the elucidation of MSC secretome roles in the context of dermal fibroblasts physiology. This secretome fractionation and functional comparison strategy using in vitro cell-based model and comparative transcriptomics analysis would be a useful tool for comprehensive functional evaluation of cellular secretome.

**Figure 1.**
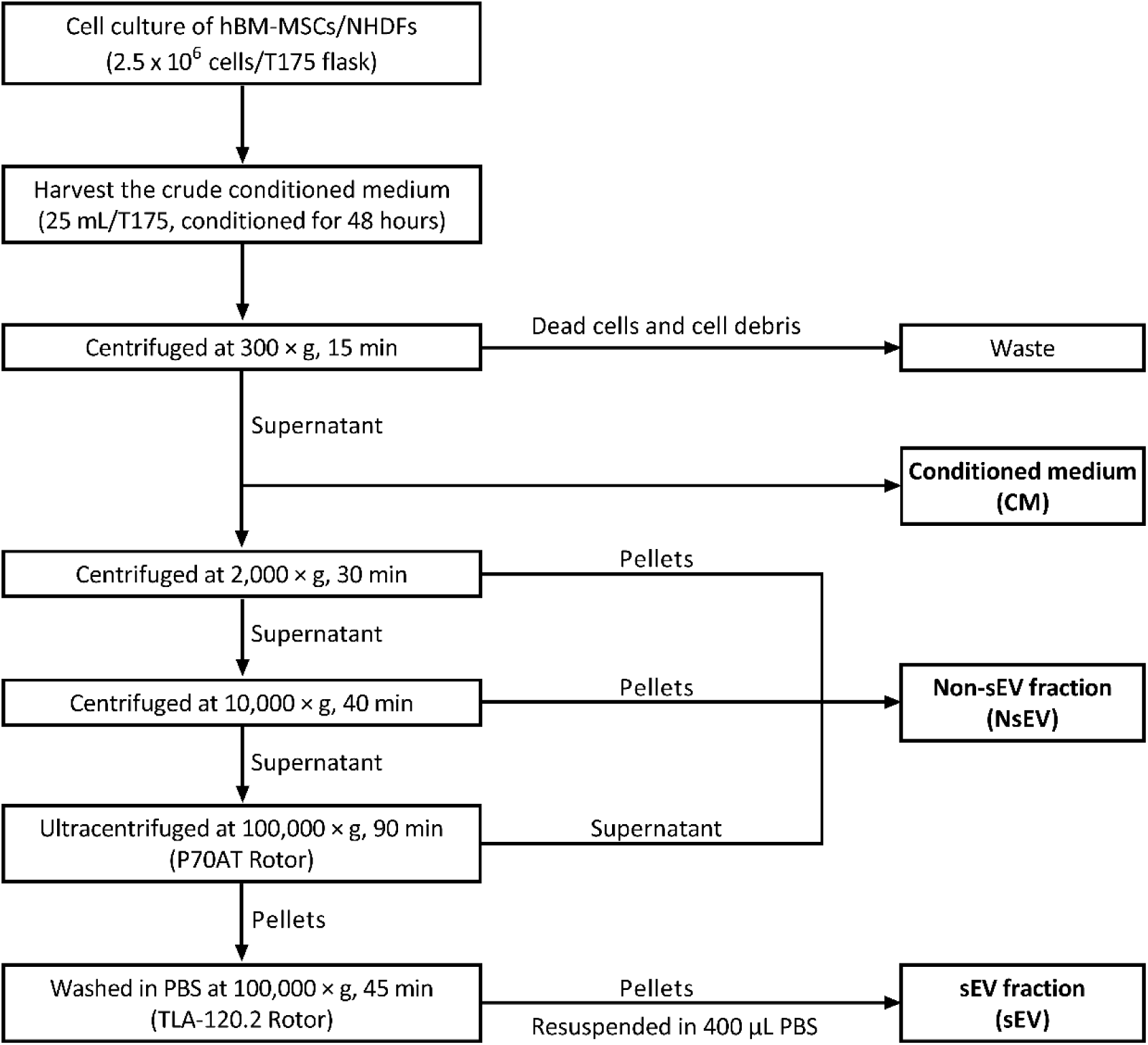
Flow chart of the fractionation protocol of secretome fractions.

## Materials and Methods

### Cell Cultivation

Human Bone Marrow derived Mesenchymal Stromal Cells (hBM-MSCs) (Cat.: C-12974, Lot.: 438Z012.1) and Normal Human Dermal Fibroblasts (NHDFs) (Cat.: C-12302, Lot.: 412Z029.2) at passage 2 (P2) were purchased from PromoCell, Germany. The hBM-MSCs were subcultured to passage 9 (P9) in mesenchymal stromal cell growth medium (Cat.: C-28009, Lot.: 466M132, PromoCell, Germany) with supplement (Cat.: C-39809, Lot.: 465M187, PromoCell, Germany) and 1% Antibiotic-Antimycotic (Cat.: 15240-062, Lot.: 2257209, Gibco, USA), and NHDFs to P10 in fibroblast growth medium (Cat.: C-23020, Lot.: 466M324, PromoCell, Germany) with supplement (Cat.: C-39325, Lot.: 468M030, PromoCell, Germany) and 1% Antibiotic-Antimycotic. Cell morphology was observed by OLYMPUS CKX53 optical microscope (OLYMPUS, Japan)

### Depletion of Extracellular Vesicle (EV) from Fetal Bovine Serum (FBS)

Fetal bovine serum derived extracellular vesicles (FBS-EV) were depleted from FBS (Cat.: 10099-141C, Lot.: 2139378CP, Gibco, USA) by ultracentrifugation at 120,000 × g, 4℃ for 18 hours using Himac CP70NE Ultracentrifuge (Himac, Japan) with P28S Rotor (Himac, Japan). The supernatant was collected and sterilized by filtering through 0.22 μm vacuum filter apparatus (Cat.: 431097, Corning, USA). The sterile EV-depleted FBS was obtained and stored at -80 ℃ before use.

### Preparation of Conditioned Medium of hBM-MSCs and NHDFs

The hBM-MSCs (P9) and NHDFs (P10) were cultured in multilayer cell culture flasks (Cat.: 353144, Corning, USA) at a seeding density of 14285 cells/cm^2^ (equal to 2.5 × 10^6^ cells per T175 flask area), using 20 mL of corresponding PromoCell complete medium per T175 flask area at 37℃ in a humidified incubator containing 5% CO_2_. After 24 hours of attachment, cells were gently washed twice with Phosphate Buffered Saline (PBS, Cat:10010-031, Lot: 2192549, Gibco, USA) and cultured with 25 mL EV-depleted medium consisting of Dulbecco’s Modified Eagle Medium (DMEM), Low Glucose, Pyruvate (Cat:11885-084, Lot: 2192435, Gibco, USA), supplemented with 1% Penicillin-Streptomycin (Cat:15140-122, Lot: 2257223, Gibco, USA) and 10% EV-depleted FBS per T175 flask area. After incubation for further 48 hours, the conditioned medium derived from hBM-MSCs and NHDFs were harvested, respectively, centrifuged at 300 × g, 4℃ for 15 min using Eppendorf 5810R Centrifuge (Eppendorf, Germany) with A-4-81 Rotor (Eppendorf, Germany) to remove the dead cells and cell debris. 10 mL of the centrifuged conditioned medium from hBM-MSCs (MSC-CM) and NHDFs (HDF-CM) were taken, respectively, and stored at 4℃ before use (applied to NHDF-based assay within 2 days). The rest of MSC-CM and HDF-CM were used to prepare sEV/non-sEV fractions.

### Preparation of sEV Fraction and Non-sEV Fraction of hBM-MSCs and NHDFs

sEVs were isolated by differential centrifugation from MSC-CM (MSC-sEV) and HDF-CM (HDF-sEV) as previously described by Thery et al ^28^ with modification. The protocol includes 2000 × g for 30 min using Eppendorf 5810R Centrifuge with A-4-81 Rotor, 10,000 × g for 40 min using with FA-45-6-30 Rotor (Eppendorf, Germany), 100,000 × g ultracentrifugation for 90 min using Himac CP70NE Ultracentrifuge (Himac, Japan) with P70AT Rotor (Himac, Japan). All the discards from MSC-CM and HDF-CM in the above three steps were collected as non-sEV fraction, respectively, known as MSC-NsEV and HDF-NsEV. The crude pellets from MSC-CM and HDF-CM after the first-round 100,000 × g ultracentrifugation were washed with PBS at 100,000 × g and 4 ℃ for 45 min using Optima™ MAX-TL Ultracentrifuge (Beckman Coulter, USA) and TLA-120.2 Rotor (Beckman Coulter, USA), respectively. The pellets after second-round ultracentrifugation from MSC-CM and HDF-CM were resuspended in 400 µL PBS, respectively, known as MSC-sEV and HDF-sEV. A flow chart of the fractionation protocol of secretome fractions was shown in Figure 1.

### Nanoparticle Tracking Analysis (NTA)

sEV samples were diluted 100-fold with PBS before measurement. The movement of the particles were recorded for 5 x 60s at camera level 13 by NanoSight NS300 (Malvern Panalytical Ltd, UK) and analysed by NTA 3.2 software (Malvern Panalytical Ltd, UK) at detect threshold 7 for EV concentrations and size distributions.

### Protein Extraction and Quantification

After the collection of conditioned medium, hBM-MSCs (P9) and NHDFs (P10) were trypsinised with 5 mL of 0.05% trypsin-EDTA solution (Cat.: 25300-054, Lot.: 2185895, Gibco, USA) per T175 flask area for 2 min, respectively, inactivated with 10 mL completed medium per T175 flask area and centrifuged at 300 × g, 20℃ for 5 min to get the cell pellets. The cell pellets of hBM-MSCs (P9) and NHDFs (P10) were lysed by adding Pierce^TM^ RIPA buffer (Cat.: 89901, Lot.: VI311555, Thermo Fisher Scientific, USA) with freshly added cOmplete™, Mini, EDTA-free Protease Inhibitor CocktailTM (Cat.: 04693159001, Lot.: 43002700, Roche, Switzerland), respectively. The mixture was incubated on ice for 30 min, and then centrifuged at 12000 rpm, 4℃ for 20 min using Eppendorf 5810R Centrifuge with FA-45-30-11 Rotor (Eppendorf, Germany), respectively. The supernatant was taken as MSC-lysate and HDF-lysate. sEV fraction, non-sEV fraction and the whole conditioned medium were lysed by adding RIPA buffer with freshly added protease inhibitor and incubating on ice for 30 min. Protein concentration of the cell lysate, sEV fraction, non-sEV fraction and the whole conditioned medium from hBM-MSCs and NHDFs were quantified using QuantiPro^TM^ BCA Assay Kit (Cat.: QPBCA-1KT, Lot.: SLCH0421, Sigma-Aldrich, USA).

### Western Blot

Cell lysate and sEV lysate of hBM-MSCs and NHDFs were normalised to the same protein concentration by PBS, based on the protein quantification results. The lysates were mixed with 4x Laemmli sample buffer (Cat.: 1610747, Lot.: 64351030, Bio-Rad, USA) and NuPAGE sample reducing agent (Cat.: NP0004, Lot.: 2205873, Thermo Fisher Scientific, USA) depend on the type of antibody, and then boiled at 95℃ for 5 min. The lysates and Precision Plus Protein Kaleidoscope Prestained Protein Standard (Cat.: 1610375, Lot.: 64313981, Bio-Rad, USA) were loaded and run on 4–20% Mini-PROTEAN® TGX™ Precast Protein Gels (Cat.: 4561094, Lot.: 64360183, Bio-Rad, USA) at a constant voltage of 120V for 65 min in a Mini-PROTEAN Tetra Cell (Cat.: 1658001, Bio-Rad, USA) filled with diluted Tris-Glycine SDS Buffer (Cat.: 1610732, Lot.: 64359578, Bio-Rad, USA), and then transferred onto a polyvinylidene fluoride (PVDF) membrane at constant voltage of 25V for 3 min using Trans-Blot TurboTransfer Packs (Cat.: 1704156, Lot.: 64340580, Bio-Rad, USA) and Trans-Blot Turbo system (Cat.: 1704155, Bio-Rad, USA). The membrane was washed in diluted 20X modified Dulbecco’s PBS Tween 20 Buffer (Cat.: 28346, Lot.: VL315361, Thermo Fisher Scientific, USA) for 5 min and then blocked with Blotting-Grade Blocker (Cat.: 1706404, Lot.: 64329693, Bio-Rad, USA) in PBS-Tween buffer at room temperature for 60 min. The blots were probed with primary antibodies at 4℃ overnight: CD63 (1:2000; Cat.: ab59479, Lot.: GR3288353-7, Abcam, UK), CD81 (1:2000; Cat.: ab79559, Lot.: GR3314758-5, Abcam, UK), calnexin (1:1000; Cat.: 2679S, Lot.: 6, Cell Signaling Technology, USA) and GM130 (1:500; Cat.: 610822, Lot.: 9140853, BD Biosciences, USA), followed by horseradish peroxidase conjugated secondary anti-mouse (1:1000; Cat.: 7076P2, Lot.: 34, Cell Signaling Technology, USA) or anti-rabbit antibody (1:1000; Cat.: 7074P2, Lot.: 28, Cell Signaling Technology, USA) incubation at room temperature for 60 min. The PVDF membranes were then exposed to Clarity Western ECL Substrate (Cat.: 170-5060, Bio-Rad, USA) for 10 min at room temperature and then exposed to Bio-Rad Chemidoc XRS system (Cat.: 1708265, Bio-Rad, USA) for imaging.

### Transmission Electron Microscopy (TEM)

sEV morphology was visualized by transmission electron microscopy. Briefly, 5 µl of solution was applied to freshly glow discharged on 300 mesh copper grids coated with a thin carbon film (Cat.: BZ10023b, ZJKY, China) for 2 min, blotted with filter paper and stained with 2% uranyl acetate (Cat.: GZ02625, ZJKY, China) for 15 s, then blotted and air dried. Grids were imaged in a Tecnai G2 F20 S-TWIN transmission electron microscope (FEI, USA) at 200 KV.

### Preparation of Different Secretome Treatments for Functional Assay

The sEVs (MSC-sEV and HDF-sEV) were diluted in fresh DMEM to concentrations of 30 μg/mL, 1 μg/mL, 0.1 μg/mL, 0.01 μg/mL, respectively. The CMs (MSC-CM and HDF-CM) and NsEVs (MSC-NsEV and HDF-NsEV) were diluted in fresh DMEM to concentrations of 1500 μg/mL, 750 μg/mL, 30 μg/mL, 1 μg/mL, respectively. As sEV pellets were suspended in PBS, an equal volume of PBS vehicle was added to fresh DMEM as DMEM control (0 μg/mL).

### sEV-uptake Assay

NHDFs at passage 8 were seeded at 4000 cell/cm^2^ with 500 µL of NHDF growth medium, making around 8000 cells per well in a Costar 24-well plate (Cat.: 3524, Corning, USA). After 24 hours of attachment, cells were gently washed once with PBS. The sEVs (MSC-sEV and HDF-sEV) were freshly labelled with PKH67 green fluorescent dye (Cat.: P7333-1ML, Lot.: MKCK0731, Sigma-Aldrich, USA) according to the manufacturer’s protocol, with some modifications. Briefly, 8 μL PKH67 was dissolved in 2 mL Diluent C (Cat.: CGLDIL-10ML, Lot.: MKCK8658, Sigma-Aldrich, USA) to prepare a 2x stain solution (4 × 10^-6^ M). Separately, sEV pellet was resuspended in 1 mL Diluent C. The same volume of PBS was mixed with 1 mL of Diluent C parallelly as a control group. The sEV suspension was mixed with an equal volume of 2x stain solution and incubated for 10 min. The labelling reaction was stopped by adding an equal volume of 1% BSA (Cat.: A1933-25G, Lot.: WXBD2613V, Sigma-Aldrich, USA) in PBS to bind excess dye. The labelled sEVs were washed once with PBS by ultracentrifugation at 100,000 ×g for 90 min. Fresh DMEM supplemented with PKH67 labelled sEVs (30 μg/mL) was used to incubate the NHDFs. The PBS control group was treated following the same protocol. After 6-hour coculture, the NHDFs were washed once with PBS, fixed with 4% paraformaldehyde (Cat.: 43368, Lot.: M21H030, Alfa Aesar, China) for 15 min, and then treated with 0.5% Triton® X-100 (Cat.: T9284-100ML, Lot.: SLBF2001V, Sigma-Aldrich, USA) for 15 min for cell membrane permeation, washed 3 times with PBS for 5 min each time between and after each step. The nuclei were stained by adding 1 μg/mL DAPI stain solution (Cat.: 62248, Lot.: VC2962251, Thermo Fisher Scientific, USA) for 2 min following the manufacturer’s protocol, washed 3 times with PBS for 5 min each time between and after each staining step. 500 µl PBS was added to each well to keep the cells hydrated while imaging. The cells were imaged by FV3000 confocal microscope (OLYMPUS, Japan).

### Proliferation Assay

NHDFs at passage 8 were seeded at 15000 cells/cm^2^ with 100 µL of NHDF growth medium, making 5000 cells per well in a Costar 96-well plates (Cat.: 3599, Corning, USA). After 24 hours of attachment, cells were gently washed once with PBS. Different secretome treatments (CMs, sEVs, NsEVs from hBM-MSCs and NHDFs) at different concentrations were used to incubate the NHDFs for 3 days, respectively. The day when the original NHDF growth medium replaced with different secretome treatments was deemed as Day 0. HDF proliferation assay was performed using the Cell Counting Kit-8 (Cat.: CK04, Lot.: PF724, Dojindo, Japan). CCK-8 reagent and DMEM was mixed at volume ratio of 1:10. On Day 0, cells were washed once with PBS and replaced with 110 µL CCK-8 and DMEM mixture in each cell testing wells and blank wells (110 µL CCK-8 and DMEM mixture without cells). Absorbance expressed as Optical Density (OD) was assessed at 450 nm (OD 450) using a Tecan Spark microplate reader (Tecan, Switzerland) after 2 hours of incubation at 37 °C. OD values were determined on Day 0, Day 1, Day 3 at the same time-point of the day using the same protocol. Triplets were performed for each secretome treatment group on each day. All the OD450 values were normalised as ratios of OD450 at Day 0 by dividing OD450 values of Day 0.

### Migration Assay

NHDFs at passage 8 were seeded at 50000 cell/cm^2^ with 1.5 mL of NHDF growth medium, making 100000 cells per well in a 24-well plate with a 2-well culture-insert (ibidi, Germany). After 24 hours of attachment, a confluent cell layer was resulted in each well and then 2-well culture inserts were gently removed by using sterile tweezers, leaving a cell-free gap of 500 μm in the middle of each well of the plate. Cells were gently washed once with PBS and different secretome treatments (CMs, sEVs, NsEVs from hBM-MSCs and NHDFs) at different concentrations were used to incubate the NHDFs in triplets for 60 hours. The time point when the original NHDF growth medium replaced with different secretome treatments was deemed as 0 h. Cell performance was recorded as bright field images by optical microscope using a 4x objective at 0 h, 6 h, 12 h, 18 h, 24 h, 30 h, 36 h, 42 h, 48 h, 60 h. The images were analysed by ImageJ and the cell covered area of the gap was quantified in μm^2^ using Matlab.

### RNA extraction and Bulk RNA-sequencing analysis

NHDFs at Passages 6-8 were seeded in 6-well plates at a seeding density of 200,000 cells per well and cultured for 24 hours in standard supplemented medium. The medium was then changed to DMEM 1% Penicillin/Streptomycin containing 30 µg/ml of the described secretome fractions, namely MSC-CM, MSC-DM, MSC-NsEV and MSC-sEV. Control was solely DMEM 1% Pen/Strep. Cells were incubated for 48 hours with secretome fractions. Total RNA was extracted from trypsin-harvested cells with the RNeasy Mini Kit (Qiagen, cat. 74104) according to the manufacture’s protocol. The purity and amount of the isolated RNA was assessed using NanoDrop One (ThermoFisher Scientific, US) to ensure high-quality RNA (A260/A280 > 2). 200-250 ng of RNA was shipped to Novogene Ltd. (Cambridge, UK) for library preparation and mRNA sequencing. Reads were mapped to a human reference genome (GRCh38/p14) and quantified using the Salmon quant tool (Galaxy Europe). Differentially expressed genes (DEGs) analysis was performed using the R DESeq2 package (version 1.42.0), which modelled the gene level read count data assuming a negative binomial distribution and employed the over dispersed Poisson model and an empirical Bayes procedure to moderate the degree of over dispersion across genes, to identify significant DEGs. DEGs were evaluated by meeting the two criteria: (i) more than twofold change in expression; (ii) adjusted p < .05. The fold change of each gene was calculated by comparing the standardised read counts of NHDFs exposed to the chosen MSC secretome fraction compared to those cultured only with DMEM 1% Penicillin/Streptomycin (fold change = standardised read counts of treatment group/control read counts). Log2 scale of fold change was used for convenience. |log2(fold change) | > 1 means at least twofold change. GEO Accession number is GSE251807 for access to sequencing data.

### Statistical Analysis

Statistical analysis of the data was performed using IBM SPSS Statistics, version 28 and GraphPad Prism, version 8. Data were expressed as mean ± standard deviations

(SD). Each experiment was repeated at least three times. Data were analysed by General Linear Model (GLM) ANOVA and Tukey method was chosen for post-hoc multiple comparisons to assess statistical significance. Probability value below 0.05 was considered statistically significant. * means P < 0.05, ** means P < 0.01, *** means P < 0.001, **** means P < 0.0001, ns means not significant. Graphs were delineated by GraphPad Prism, version 8.

## Results

### Characterisation of MSC-sEV and HDF-sEV

sEV samples were characterised in terms of size, surface marker expressions, morphology and yield. Both MSC-sEV and HDF-sEV displayed a bell-shaped curve with a peak at 118.87 nm and 117.60 nm, and mean particle diameters at 143.60 nm and 144.07 nm, respectively, consistent with characteristic size range of sEVs (Figure 2a-c). No significant difference was found in size distribution or mean particle size between MSC-sEV and HDF-sEV. Both MSC-sEV and HDF-sEV were positive for typical EV markers such as CD63 and CD81, and negative for intracellular compartment markers like Calnexin and GM130, which confirmed the purity of the sEV samples (Figure2d). Classic cup-shaped sEV particles were found in TEM images (Figure 2e). The yield of sEVs was quantified as EV-associated protein amount by BCA method, showing that 45.27 μg for MSC-sEV and 40.68 μg for HDF-sEV can be obtained from 100 mL corresponding conditioned medium, respectively (Figure 2c).

**Figure 2.**
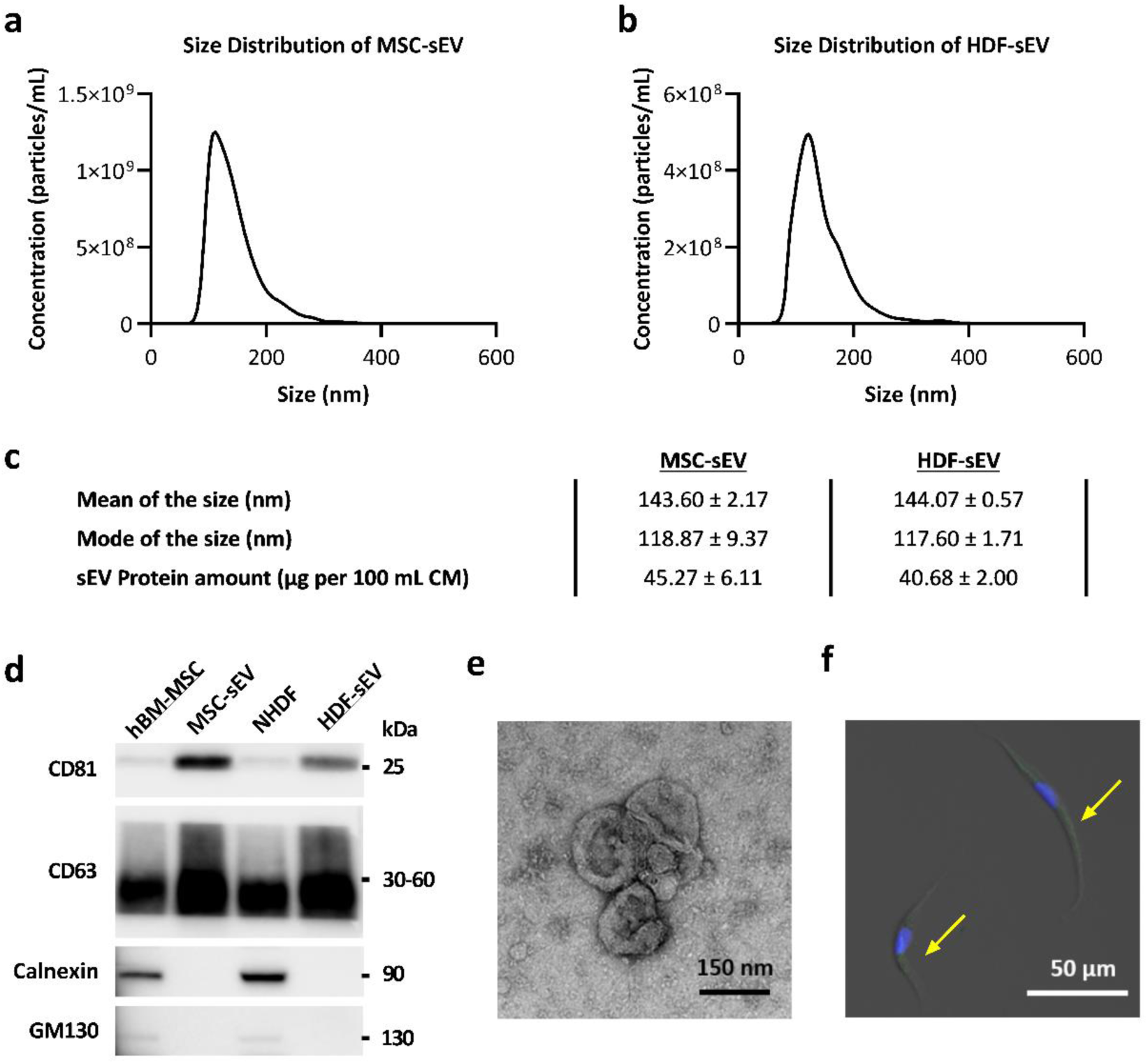
Characterisation of MSC-sEV and HDF-sEV. (a) Average size distribution of MSC-sEV samples determined by NTA (n=15). (b) Average size distribution of HDF-sEV samples determined by NTA (n=15). (c) Size (n=3) and yield (n=3) of MSC-sEV and HDF-sEV. (d) Western blot analysis of MSC-sEV, HDF-sEV and their parental cell lysates. Original blots/gels are presented in Supplementary Figure 1. (e) TEM image of MSC-sEVs. (f) Fluorescence images of MSC-sEVs taken by NHDFs. Cell nucleus of NHDFs were stained by DAPI. MSC-sEVs were stained by PKH67 (green signal was pointed out by arrows).

### Internalisation of sEVs by NHDFs

To determine whether the recipient cell NHDFs can internalise sEVs, PKH67-labelled MSC-sEVs were incubated with NHDFs for 6 hours. The cellular uptake of MSC-sEVs by NHDFs was confirmed by the confocal image. (Figure 2f).

### The Effect of Cellular Secretome on Migration of NHDFs

To determine the effects of different secretome treatments on migration of NHDFs, the migration assay was performed on NHDFs treated with MSC-CM, MSC-sEV, MSC-NsEV at different concentrations (0 μg/mL, 1 μg/mL, 30 μg/mL) for 60 hours. Blank (DMEM) control group was represented as 0 μg/mL. Representative bright field images of NHDFs migration were shown in Supplementary Figure 2. To compare the difference in a quantitative way, all the bright field images were processed by ImageJ and the cell covered area of the gap changed with time course was quantified by Matlab. General Linear Model (GLM) ANOVA was used to analyse factors of migration time, secretome fraction type and concentration of secretome. As expected, migration time (Hour) [F(9, 756) = 2032.78, p < 0.001] was a variable that significantly affected the migration of NHDFs (see Supplementary Table 1). A typical growth curve of migration consist of a linear growth phase and a saturation phase was showed in Figure 3a, indicating the migration velocity was nearly constant when cell migration started and then slowed down to zero when the NHDFs reached confluency at 42 h, probably due to lack of space. Migration of NHDFs was also significantly affected by the secretome fraction type (Treatment) [F(2, 756) = 29.77, P < 0.001] and concentration of secretome (Concentration) [F(2, 756) = 11.48, P < 0.001], and significant interaction was observed between these two variables [F(4, 756) = 11.38, P < 0.001], showing different functional dynamics of the components in MSC-CM, MSC-sEV and MSC-NsEV (see Supplementary Table 1). Tukey’s post-hoc multiple comparisons were therefore performed between different secretome fractions of MSCs at different concentrations for their effects on migration behaviour of NHDFs (see Supplementary Table 2) and an interaction plot (Figure 3b) was showed based on the results of Tukey’s post-hoc multiple comparisons (the growth curve of migration of each group was expressed by its mean on the interaction plot). Generally, both MSC-CM and MSC-sEV mediated an overall positive effect on migration of NHDFs when compared to blank control group (0 μg/mL), and it was shown that the performance line of MSC-sEV was consistently above that of the MSC-CM (Figure 3b). Specifically, MSC-sEV started to show promotive effect on NHDF migration at lower dose (1 μg/mL) than MSC-CM (30 μg/mL), although no significance difference was found between them at higher dose (30 μg/mL). There was no significant difference found between MSC-NsEV treatments at dose ≤30 μg/mL and blank control group.

**Figure 3.**
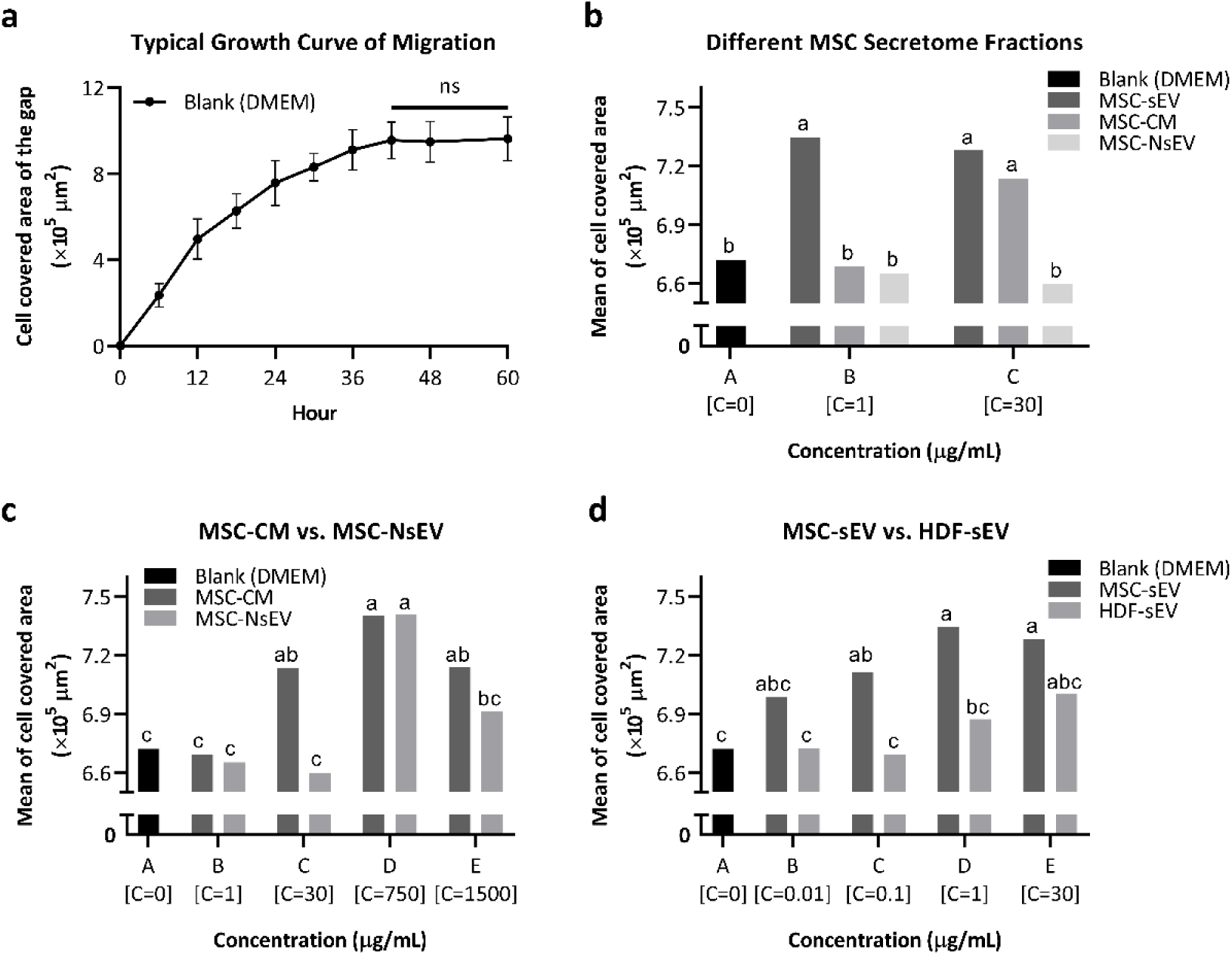
The effect of cellular secretome at different concentration (represented as C) on migration of NHDFs. (a) A typical growth curve of migration. NHDFs in all treatment groups showed growth curves in a similar trend. The migration curve of NHDFs treated with blank (DMEM) was shown here. (b) Interaction plot for migration behaviour of NHDFs in response to factors of secretome fraction type from MSCs (MSC-sEV, MSC-CM and MSC-NsEV) and concentration of secretome based on the results of Tukey’s post-hoc multiple comparisons (see raw data in Supplementary Figure 3a&b). (c) Interaction plot for migration behaviour of NHDFs in response to factors of secretome fraction type from MSCs (MSC-CM and MSC-NsEV) and concentration of secretome (including high concentrations) based on the results of Tukey’s post-hoc multiple comparisons (see raw data in Supplementary Figure 3c-f). (d) Interaction plot for migration behaviour of NHDFs in response to factors of sEV types (MSC-sEV and HDF-sEV) and concentration of sEVs based on the results of Tukey’s post-hoc multiple comparisons (see raw data in Supplementary Figure 3g-j). P < 0.05 was deemed as a statistically significant difference. ns means not significant in (a). Means that do not share a letter are significantly different in (b), (c) and (d).

To further investigate the difference of the performance between MSC-CM and MSC-NsEV on NHDF migration, NHDFs were treated with MSC-CM and MSC-NsEV at high concentrations (750 μg/mL, 1500 μg/mL). Again, all the images were processed and quantified by ImageJ and Matlab to compare the difference in a quantitative way. GLM ANOVA and Tukey’s post-hoc multiple comparisons were performed (see Supplementary Table 3 and Supplementary Table 4) and the interaction plot (Figure 3c) showed that both two types of secretome enhanced the NHDF migration ability at high concentration treatment, compared to the blank control group. Particularly, MSC-CM started to show a significance promotive effect on NHDF migration at 30 μg/mL, and the level of enhancement firstly increased and then showed a reduction with increased dose level, suggesting that there might be both synergic and antagonistic components in the CM, and there might be a critical value as optimal dose for NHDF migration in the range of 30 – 1500 μg/mL, although no significant difference was found between 30 μg/mL, 750 μg/mL and 1500 μg/mL. A similar trend was demonstrated with increased dose of MSC-NsEV treatments, while significant improvement on NHDF migration was only observed at 750 μg/mL among different concentration treatments, compared with blank control group.

Based on the previous results, evidence suggested that sEV fraction might display superior ability than NsEV fraction for NHDF migration. Therefore, to further investigate the effect of sEV fraction, NHDFs were treated with sEVs from different cell sources (MSC-sEV and HDF-sEV) at several concentrations (0 μg/mL, 0.01 μg/mL, 0.1 μg/mL, 1 μg/mL, 30 μg/mL) and the migration performance was recorded for 60 hours. GLM analysis was performed (see Supplementary Table 5), indicating that cell source of sEVs (sEV type) [F(1, 836) = 30.40, P < 0.001] was a variable that significantly affected NHDF migration ability, and it significantly interacted with concentration of sEVs (Concentration) [F(4, 836) = 2.49, P = 0.042], showing different functional dynamics of the MSC-sEV and HDF-sEV. Tukey’s post-hoc multiple comparisons were performed (see Supplementary Table 6) and an interaction plot (Figure 3d) was showed. The performance line of MSC-sEV treated NHDFs was consistently above that of HDF-sEV in Figure 3d, showing that MSC-sEV mediated an overall promotive effect on NHDF migration when compared to the sEVs produced by NHDFs itself (HDF-sEV). Especially, a dose-dependent trend of MSC-sEV was observed for NHDF migration as the level of enhancement firstly increased and then reached a plateau with increased dose level. MSC-sEV started to show a significance promotive effect on NHDF migration when the dose level ≥ 0.1 μg/mL, although no significant difference of enhancement level was found between the concentration treatments ≥ 0.1 μg/mL. However, HDF-sEV at dose range 0 – 30 μg/mL didn’t show any significant positive effect on the migration of NHDFs compared to blank control group, although a similar dose-dependent trend probably existed.

### The Effect of Cellular Secretome on Proliferation of NHDFs

To investigate the effects of different cellular secretome treatments on proliferation of NHDFs, Cell Counting Kit-8 assay was performed on NHDFs and absorbance expressed as Optical Density (OD) was assessed at 450 nm (OD 450). All the OD450 values were normalised as folds of OD450 at Day 0 by dividing OD450 values of Day 0. To compare the cell proliferation behaviour of NHDFs under treatments of CMs, sEVs and NsEVs from both hBM-MSCs and NHDFs at different doses (concentration of treatments ≤ 30 μg/mL were considered as low doses, 750 μg/mL and 1500 μg/mL were considered as high doses), factors of treatment time, cell source of secretome and secretome fraction type were evaluated by GLM (see Supplementary Table 7). As expected, the CCK8 results were significantly affected by the factor of treatment time (Day) [F(2, 526) = 407.25, p < 0.001]. A typical growth curve of proliferation was showed in Figure 4a, indicating that the NHDF proliferation behaviour represented by the folds of OD450 at Day 0 increased from Day0 to Day3 in all treatments. Out of surprise, compared to blank (DMEM) control group, all the treatments at low doses did not make any significant difference on the proliferation of NHDFs, and there was no significant difference between either different secretome fractions from the same cell source or corresponding secretome fraction from different cell sources at low doses (Figure 4b). However, with the increase of concentration, both MSC-CM and MSC-NsEV at high doses displayed significant promotive ability on the proliferation of NHDFs, indicating that there were beneficial factors for proliferation in both MSC-CM and MSC-NsEV.

**Figure 4.**
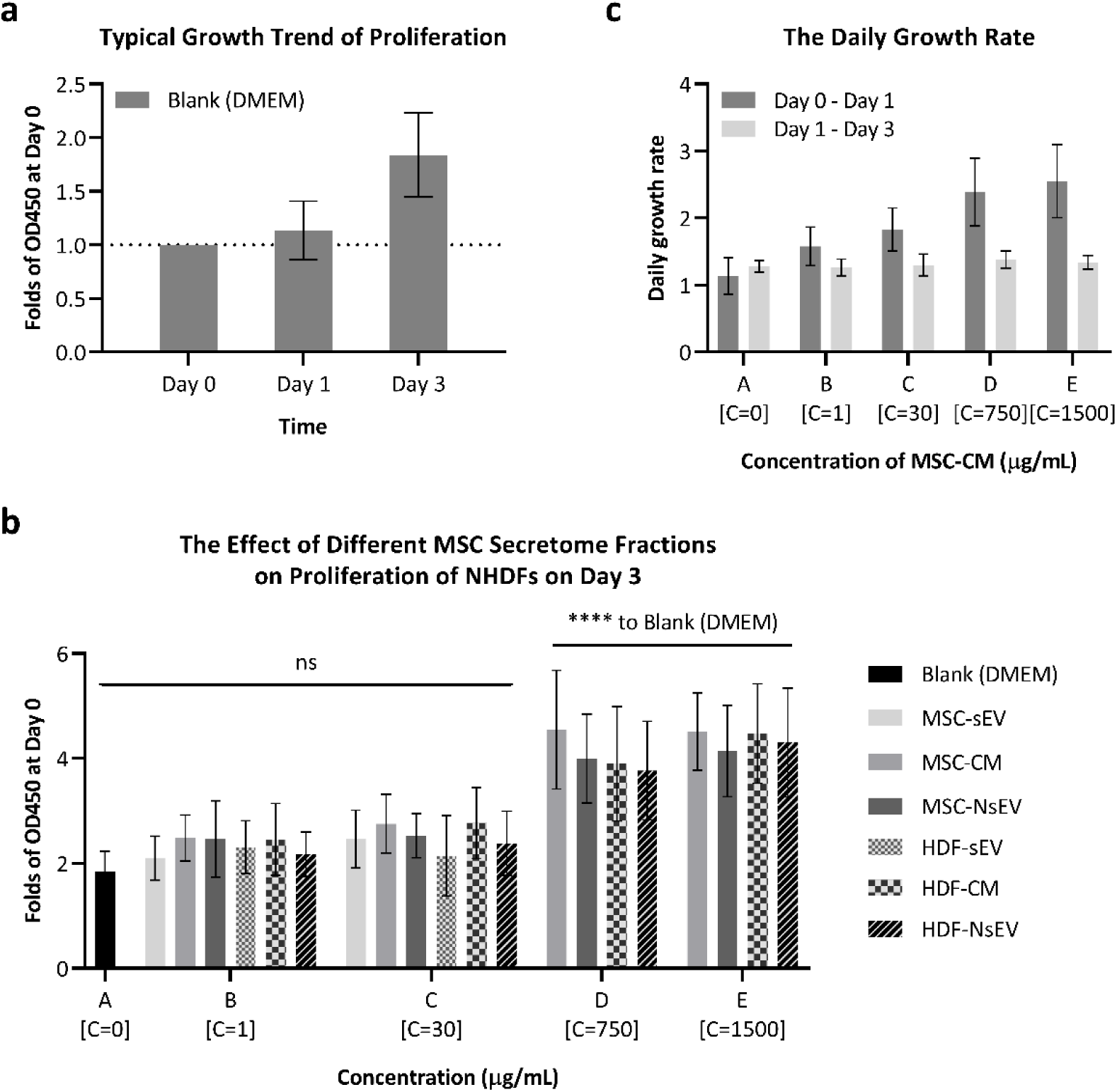
The effect of cellular secretome at different concentration (represented as C) on proliferation of NHDFs. (a) A typical growth curve of proliferation. NHDFs in all treatment groups showed growth curves in a similar trend. The proliferation curve of NHDFs treated with blank (DMEM) was chosen. (b) Plot for proliferation (Folds of OD450 at Day 0) of NHDFs at Day 3 in response to different secretome fraction types from both hBM-MSCs and NHDFs at different concentrations. Blank (DMEM) control group was represented by 0 μg/mL. P < 0.05 was considered statistically significant to Blank (DMEM) control group (see statistical analysis results in Supplementary Table 8). * means P < 0.05, ** means P < 0.01, *** means P < 0.001, **** means P < 0.0001, ns means not significant. (c) The daily growth rate during different time periods of MSC-CM treatment at different concentrations.

As it mentioned before, the NHDF proliferation behaviour represented by the folds of OD450 at Day 0 was significantly affected by treatment time, showing a clear increase trend from Day0 to Day3 in all treatments, however, the exact effective acting time for the secretome treatments was unknown. Therefore, the daily growth rate of OD450 during different time periods of MSC-CM treatment was calculated by equation (1) and equation (2):

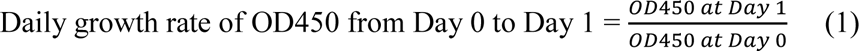

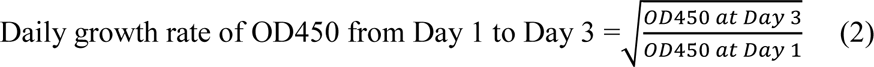

It was interesting to observe that the line of the daily growth rate of Day 0 – Day 1 were consistently above that of Day 1 – Day 3 (Figure 4c), indicating that the exact effective acting time is highly likely to be the first 24 hours after secretome treatments. Moreover, the daily growth rate of Day 1 – Day 3 remained constant, while a dose-dependent trend was found for the daily growth rate of first 24 hours, which also supports the previous result that there might be beneficial factors for proliferation in MSC-CM.

### RNA sequencing Analysis

Given that MSC-sEVs have shown to increase migration on NHDFs, but did not change proliferation, we performed RNA sequencing to determine whether exposure to MSC secretome, in particular MSC-sEVs given the migration findings, could change pathways relating to migration and proliferation. A total of 32 samples of NHDFs were sequenced with at least four biological replicates for each treatment group, namely MSC-depleted medium (DM), MSC-CM, MSC-NsEVs and MSC-sEVs, besides samples exposed only to DMEM as controls. Samples were analysed in two batches and generated comparable count distributions, though Principal Component Analysis (PCA) revealed a batch effect (Supplementary Figure 4) which was taken into consideration in the design of DESeq2 analysis and did not impact the reliability of results, given that each batch contained both controls and test samples.

A total of 1159 genes were differentially expressed when comparing controls and cells exposed to MSC-sEVs, compared to 13961 not statistically changed; the volcano plot shows the distribution of genes in regard to their significance (Figure 5a). To verify which genes had the lowest p-values, a heatmap was drawn with the 30 most differentially expressed genes (Figure 5b), which showed that samples tended to congregate according to the treatment to which they were exposed, a feature suggestive of the biological significance of different fractions. MSC-NsEV and MSC-sEV groups showed visible contrast in gene expression in comparison to the DMEM group. Some of the genes displayed are linked to proliferation and migration, such as STAT3, STC1 and DDX5.

**Figure 5.**
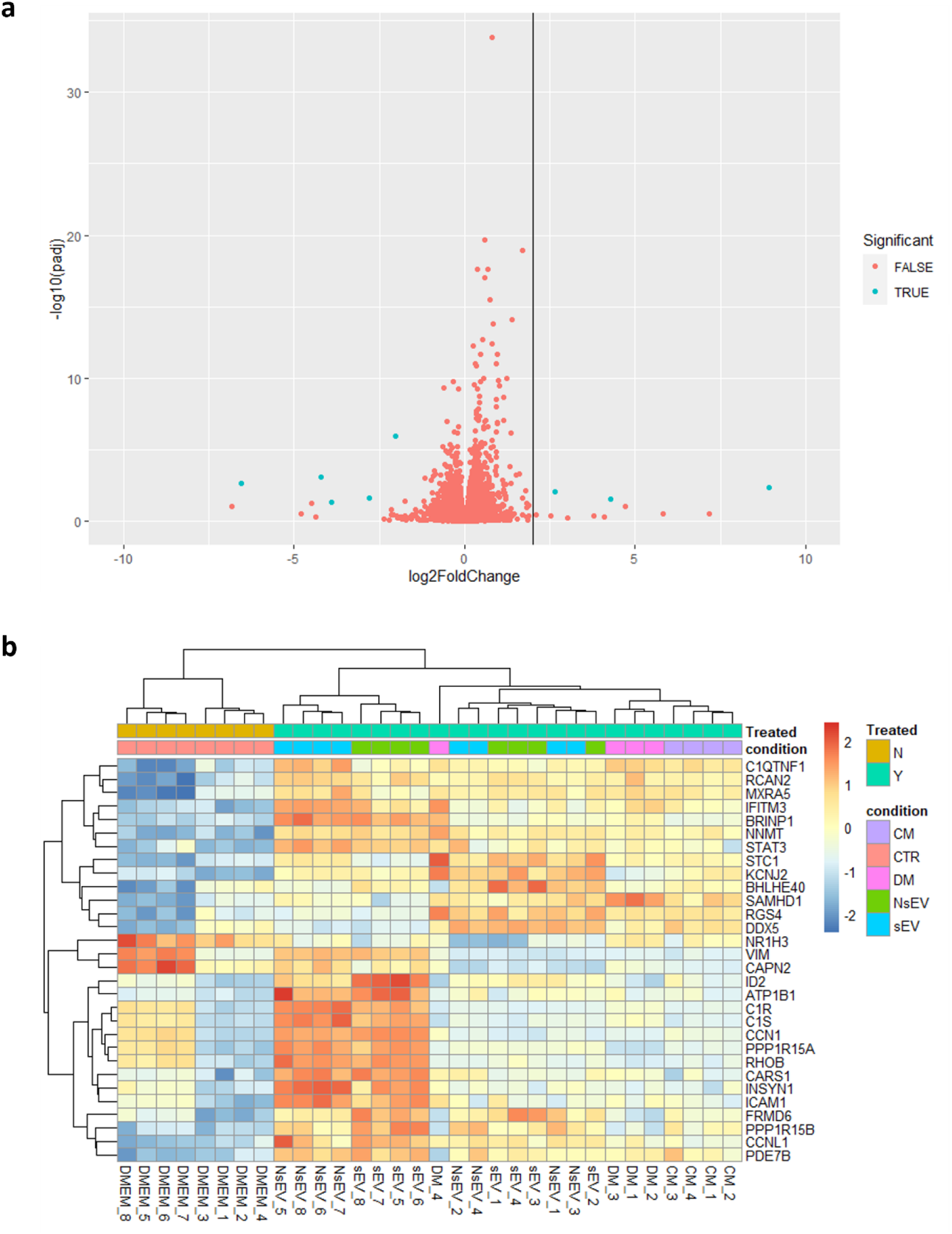
Differentially Expressed Genes (DEGs) Analysis. (a) Volcano Plot displaying proportion of DEGs. (b) Heatmap allowing comparison of expression of top 30 DEGs from each sample, which includes hierarchical cluster analysis of these DEGs between each sample. Samples are listed on the primary *x*-axis while top 30 DEGs are listed on the *y*-axis with their expression ratio (log2) expressed by the colour gradient shown. The secondary *x*-axis groups the results based on the similarity among the samples. Gene clustering is presented on the secondary *y*-axis.

### Gene Ontology

Gene Ontology analysis was performed to bring clarity to the list of DEGs we have obtained. Our purpose was to determine which pathways were significantly more represented when comparing MSC-sEV and DMEM groups, based on the DEGs between these groups. Among the most enriched pathways, several are related to migration (Figure 6a), such as epithelial cell migration, tissue migration and endothelial cell migration. Interestingly, there are many similarities in enriched pathways when we perform the analysis comparing samples exposed to MSC-NsEVs versus controls (Supplementary Figure 5). These pathways are linked, as showed in Figure 6b, and this enrichment suggest that MSC-sEVs are upregulating many genes related to migration, but there are also enriched pathways related to miRNAs biology, hinting at their importance concerning MSC-sEVs mechanisms of action. In order to visualise the interactions and overlaps of genes between enriched pathways, an UpSet plot has been used (Figure 7a), with the number of DEGs within pathways represented as bars and commonalities between pathways depicted as black lines below the bars.

**Figure 6.**
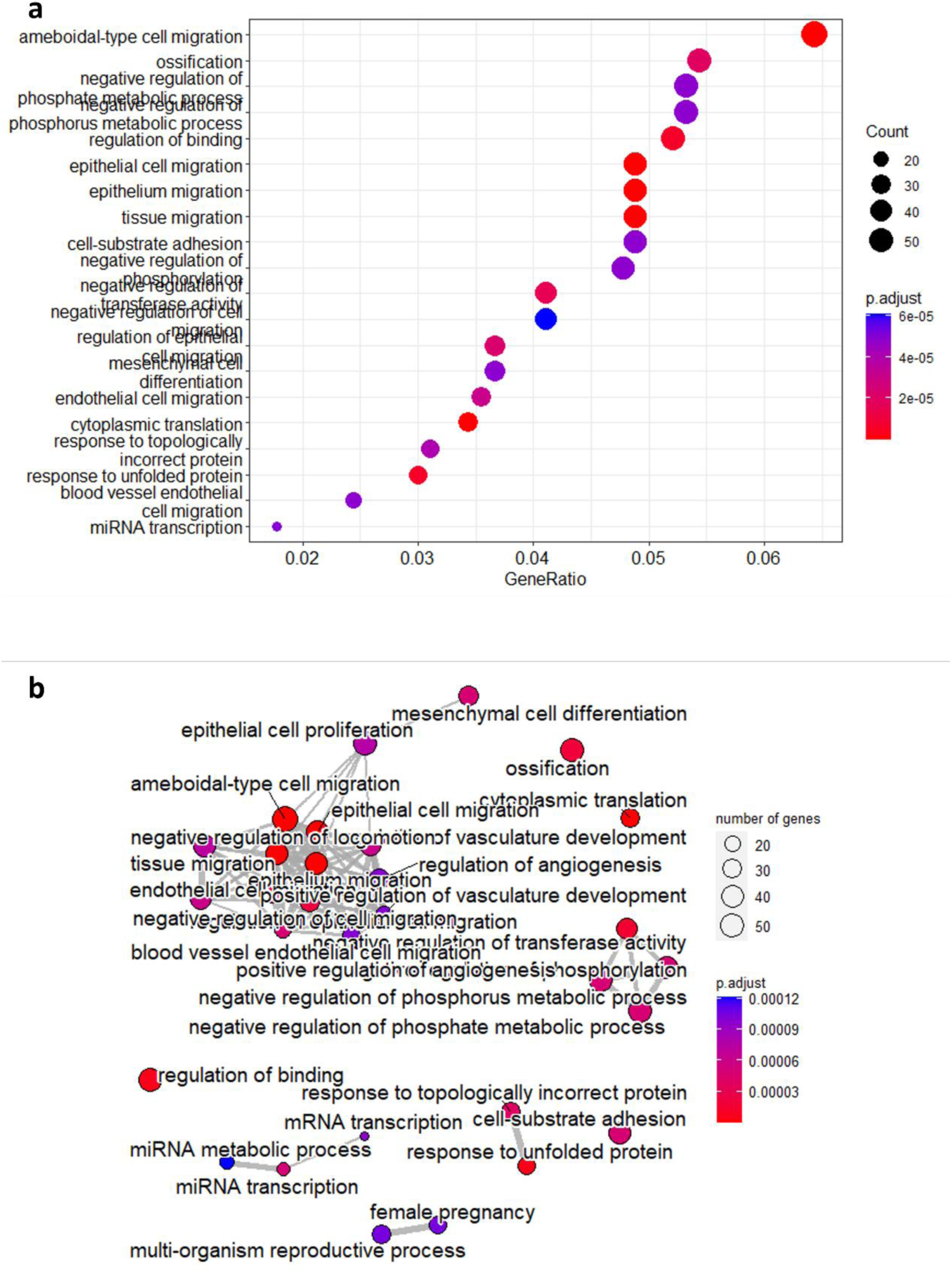
Gene Ontology Analysis and Enriched Pathways Clustering when comparing samples exposed to sEVs and controls. (a) Pathways that have been significantly changed based on the number of genes within them that are differentially expressed. (b) Links between enriched pathways and how they cluster depending on common genes.

**Figure 7.**
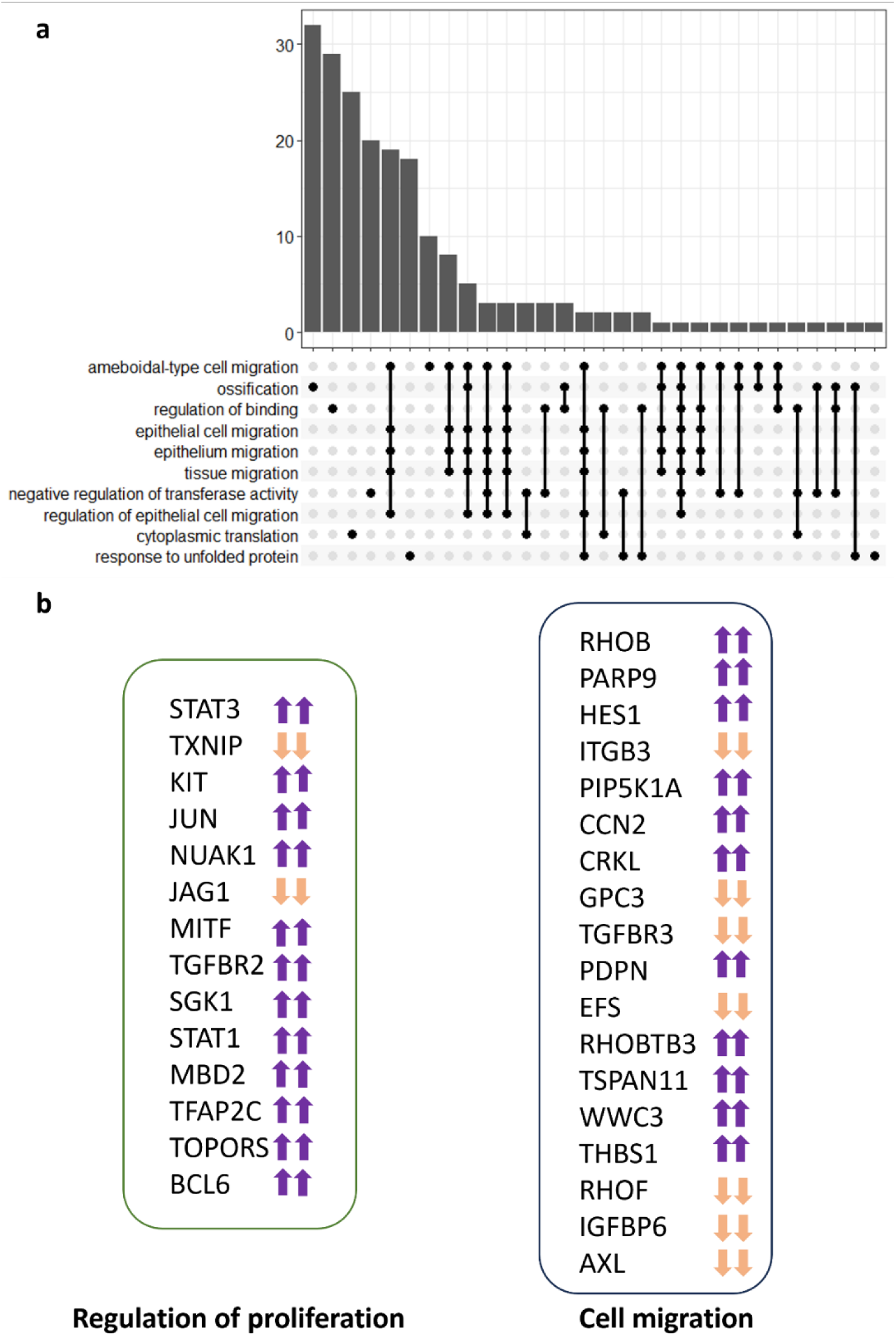
Enriched pathways links. (a) UpSet plot with bars representing the number of DEGs within the pathways enlisted and black lines denoting common genes between pathways described. (b) Differences in expression of genes that exhibited significant up-regulation (up arrow) and down-regulation (down arrow) in “Regulation of cell population proliferation” and “Cell migration” pathways. The presence of two arrows is to indicate that there is concordance between sEV and NsEV results. Cut-off p adjusted value was 0.05.

As the focus of this study was to determine the impact of MSC secretome in migration and proliferation as proxies of wound healing potential, a list of genes that have been statistically changed within the regulation of cell population proliferation (GO:0042127) and cellular migration pathways (GO:0016477) is given in Figure 7b, demonstrating that despite the lack of effect in the proliferation assay, MSC-sEVs and MSC-NsEVs have changed gene expression in a targeted manner related both to migration and proliferation with gene expression shifting in the same direction for both fractions.

## Discussion

Evidence is mounting that MSCs play various therapeutic effects through secretion and regulation of a variety of paracrine factors ^5,6^. Cellular secretome is actually a combination of membrane-bound extracellular vesicles such as sEVs, secreted soluble proteins including cytokines, chemokines, growth factors, proteases and probably some metabolism waste inside the conditioned medium. It has been verified that different components of the secretome derived from MSCs, especially MSC derived sEVs, displayed promising regenerative effects on wound healing, skin regeneration and anti-_aging 11,16,17._

According to our results, both MSC-sEV and MSC-NsEV fractions as well as MSC-CM itself showed an overall promotive effect on migration behaviour of NHDF; it is therefore reasonable to conclude that both sEV fraction and non-sEV fraction of the conditioned medium from MSCs have beneficial factors and our RNA sequencing analysis confirmed that multiple players involved in cell migration are changed in response to exposure to MSC secretome. This result is in line with previous reports that the whole conditioned medium derived from BM-MSCs significantly enhanced wound closure by accelerating the migration of keratinocytes, fibroblasts, endothelial cells, with identified wound healing mediators from MSC-CM, including TGF-β1, the chemokines IL-6, IL-8, collagen type I, fibronectin ^18–20^. Shabbir et al. and Hoang et al. claimed that sEVs derived from BM-MSCs could also display promotive effects on the migration of dermal fibroblasts by activating important signalling pathways (Akt, ERK, and STAT3) in wound healing ^14^. Indeed, our transcriptomic analysis revealed changes in the expression of genes involved in migration, including STAT1, STAT3 and TGF-β receptors, as well as enrichment of pathways responsible for epithelial and endothelial migration, promoted by MSC-sEVs.

Based on the results of the migration assay, evidence shows that MSC-sEVs are more potent in comparison to other fractions, as they showed a promotive effect at lower doses. A similar superior therapeutic ability of sEV derived from BM-MSCs were also reported by Takeuchi et al. but in promoting bone regeneration and angiogenesis, showing that MSC-sEV enhanced cellular migration, osteogenesis and angiogenesis-related gene expression, compared to MSC-CM group ^21^. MSC-sEV also mediated a significant beneficial effect on migration over sEVs produced by recipient NHDFs itself. This result might be explained by the difference in the potential of its parental cells (BM-MSCs) to accelerate wound healing, which is supported by previous studies that BM-MSCs showed greater activity than dermal fibroblasts in terms of granulation tissue formation, collagen synthesis, epithelialization and angiogenesis in vivo ^17,22–24^. Chen et al. also demonstrated that conditioned medium of BM-MSCs significantly enhanced migration and proliferation of keratinocytes and endothelial cells compared to HDF-CM ^8^. sEVs are shown to carry cell-specific cargoes such as proteins, lipids and genetic materials ^25^, served as cell-to-cell communication mediators to regulate the properties of target cells. Therefore, it is reasonable to infer that MSC-sEV may carry biological information from BM-MSCs and regulate the wound healing behaviour of the recipient NHDFs. Indeed, Hu et al. showed that MSC-sEV promoted a faster rate of wound recovery by significantly enhancing the proliferation and migration of NHDFs, and displayed amelioration of skin photoaging, compared with HDF-sEV ^26^.

There are reports on the subject of EV cargo and how it changes according to the cell type and microenvironment of parental cells, besides methodological approaches, such as different media and methods of EV isolation ^27^. Although we did not investigate miRNA cargo of sEVs in our study, our gene ontology analysis showed pathways linked to miRNA biology, which is in line with studies showing that miRNAs are key in promoting EV-mediated gene expression change in diverse pathways ^28–30^, and indeed might have been the enactors of the observed changes.

Our findings are in line with reports stating that MSC-sEV can accelerate wound closure in a dose-dependent manner ^14,31–33^, as there was a degree of dose-dependence observed in our assays. Our study also corroborates that MSCs enhance cellular proliferation and migration, which was previously reported ^34^, but showing that these cells can do so via paracrine factors through gene expression changes in several pathways even in cells that are not defective in migration.

It should be valued that the non-sEV fraction from MSCs also showed its promotive effect in NHDF migration, which might be due to the fact that the non-sEV fraction was actually the left-over of conditioned medium after isolation of sEVs, which contains those non-sEV beneficial factors including cytokines, chemokines, growth factors. However, the defective components such as some metabolism waste in the non-sEV fraction should be treated with caution.

In conclusion, our study found that different fractions of MSC derived secretome are effective in promoting migration of normal dermal fibroblasts with correspondent changes in the expression of genes within migration pathways. This study has shown that non-sEVs also have an impact in both behaviour and transcriptomics. Despite the lack of efficacy in enhancing fibroblast proliferation in cell assay, several proliferation regulatory genes have been changed after exposure to MSC-sEVs and –non-sEVs. Our study provides referential significance for the choice of cellular secretome fractions to be used in wound healing studies and trials. The secretome fractionation and functional comparison strategy using cell-based model and comparative transcriptomics analysis would be a useful tool for comprehensive functional evaluation of cellular secretome.

## Supplementary Information

**Supplementary Figure 1.**
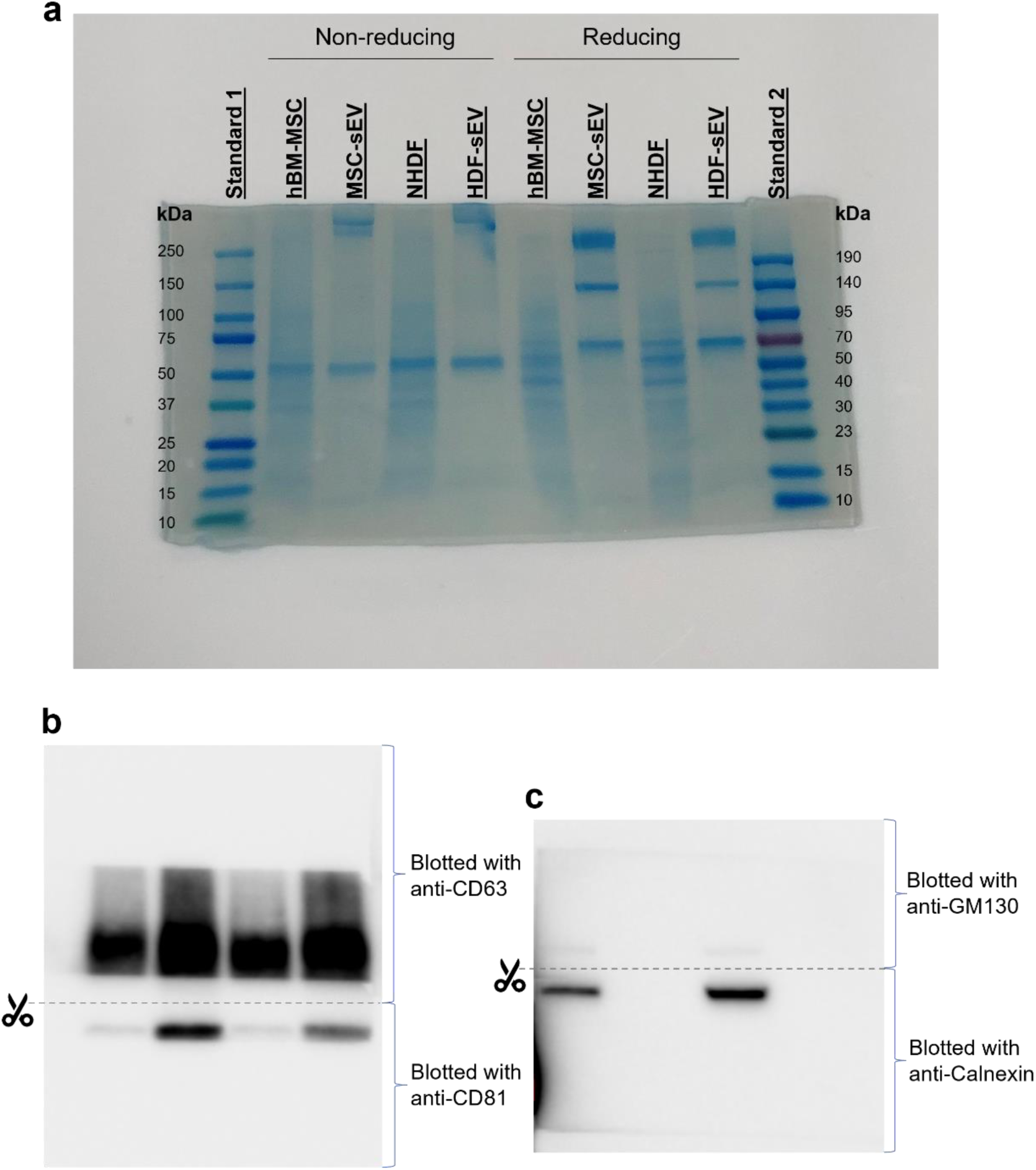
Original images from the western blot data shown in **Error! Reference source not found.**Figure 2d. (a) Proteins in the gel visualised by Coomassie Blue. Anti-CD63 antibody and anti-CD81 antibody should be used under non-reducing condition (reducing agent was not added to the sample). Anti-GM130 antibody and anti-Calnexin antibody should be used under reducing condition. (b) Western blot for CD63 and CD81. (C) Western blot for GM130 and Calnexin.

**Supplementary Figure 2.**
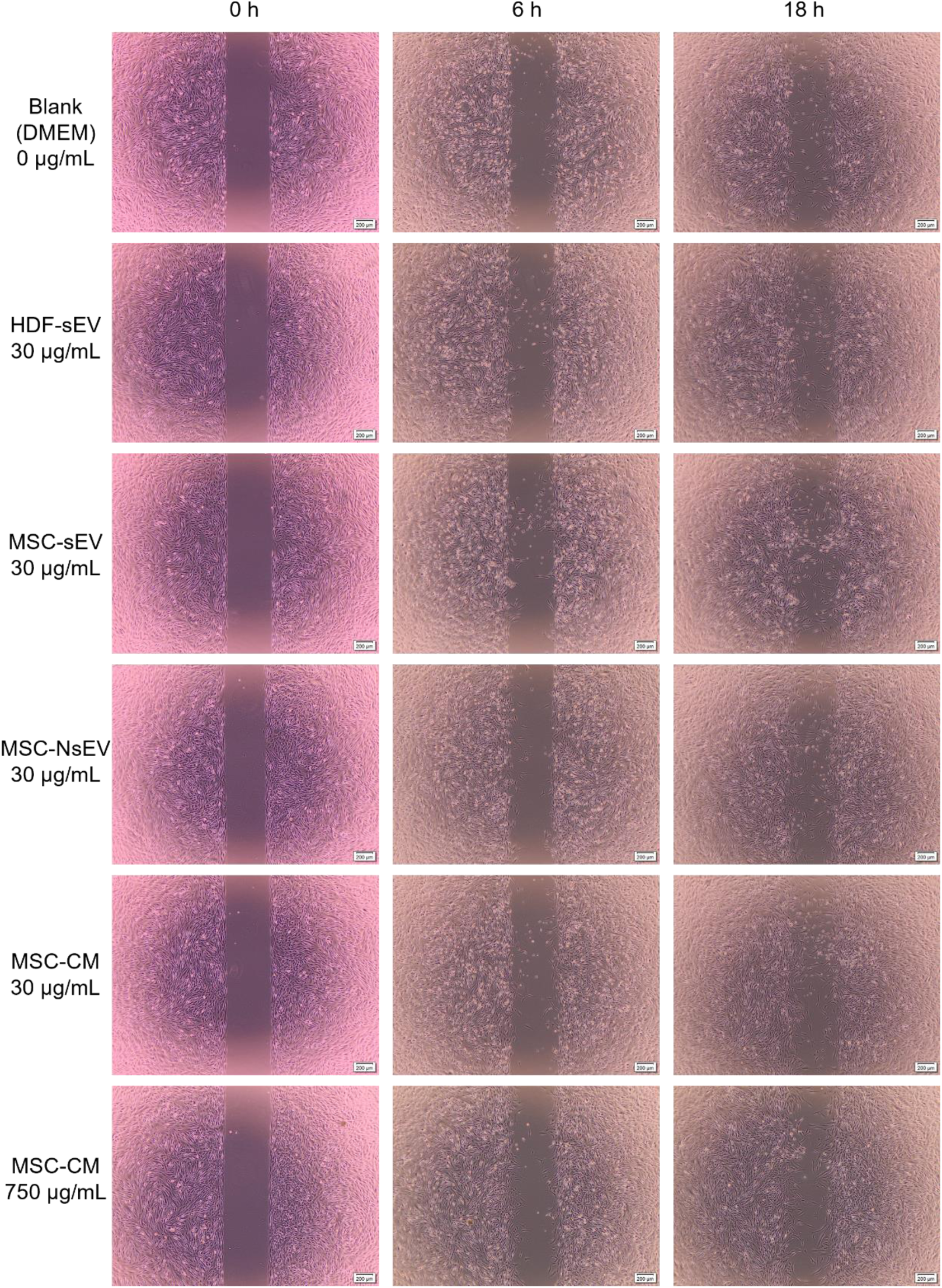
Bright field images of NHDFs treated with different secretome fractions at 0 h, 6 h and 18 h.

**Supplementary Figure 3.**
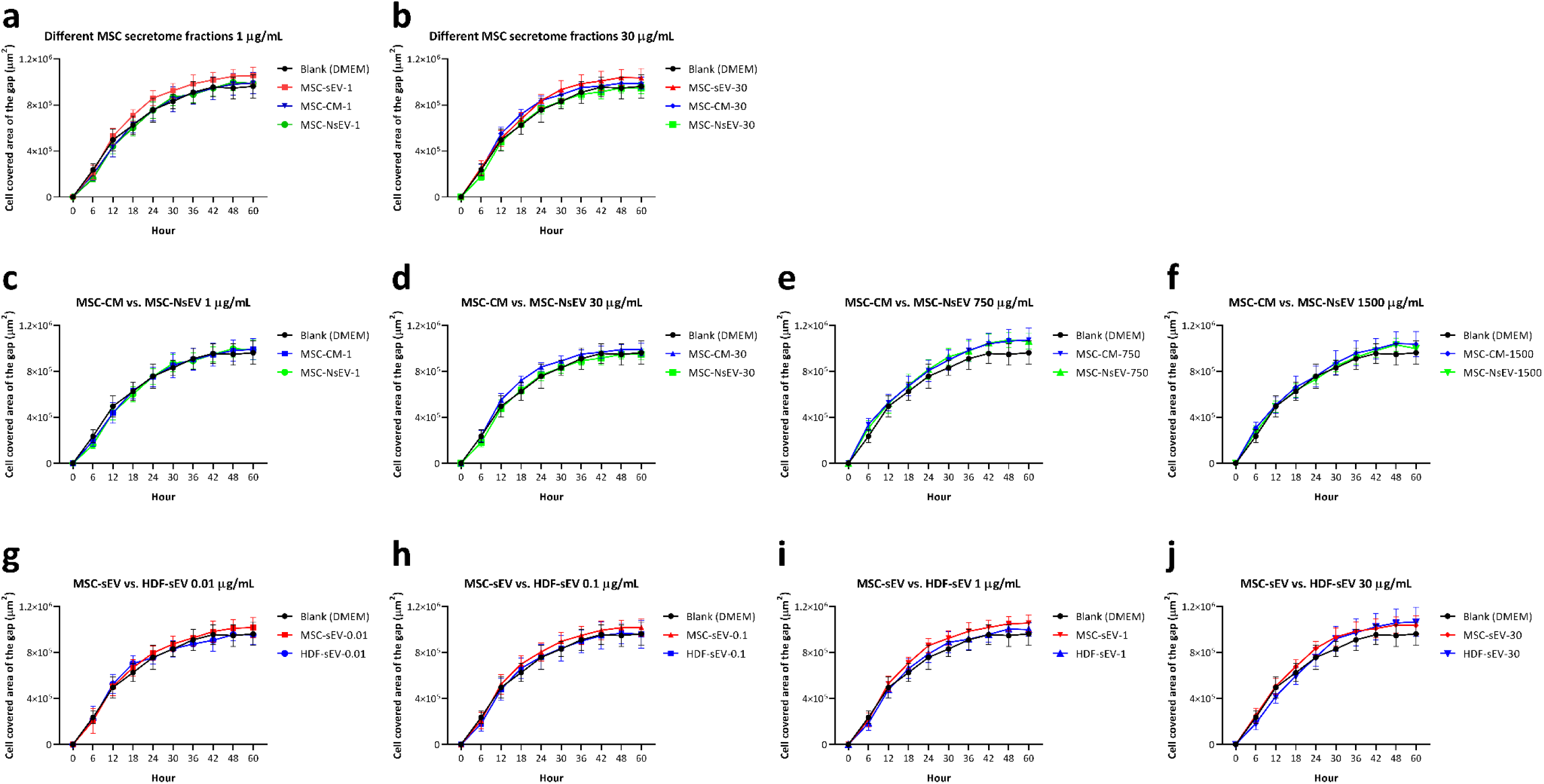
The migration curves of the NHDFs treated by different secretome fractions at different concentrations

**Supplementary Figure 4.**
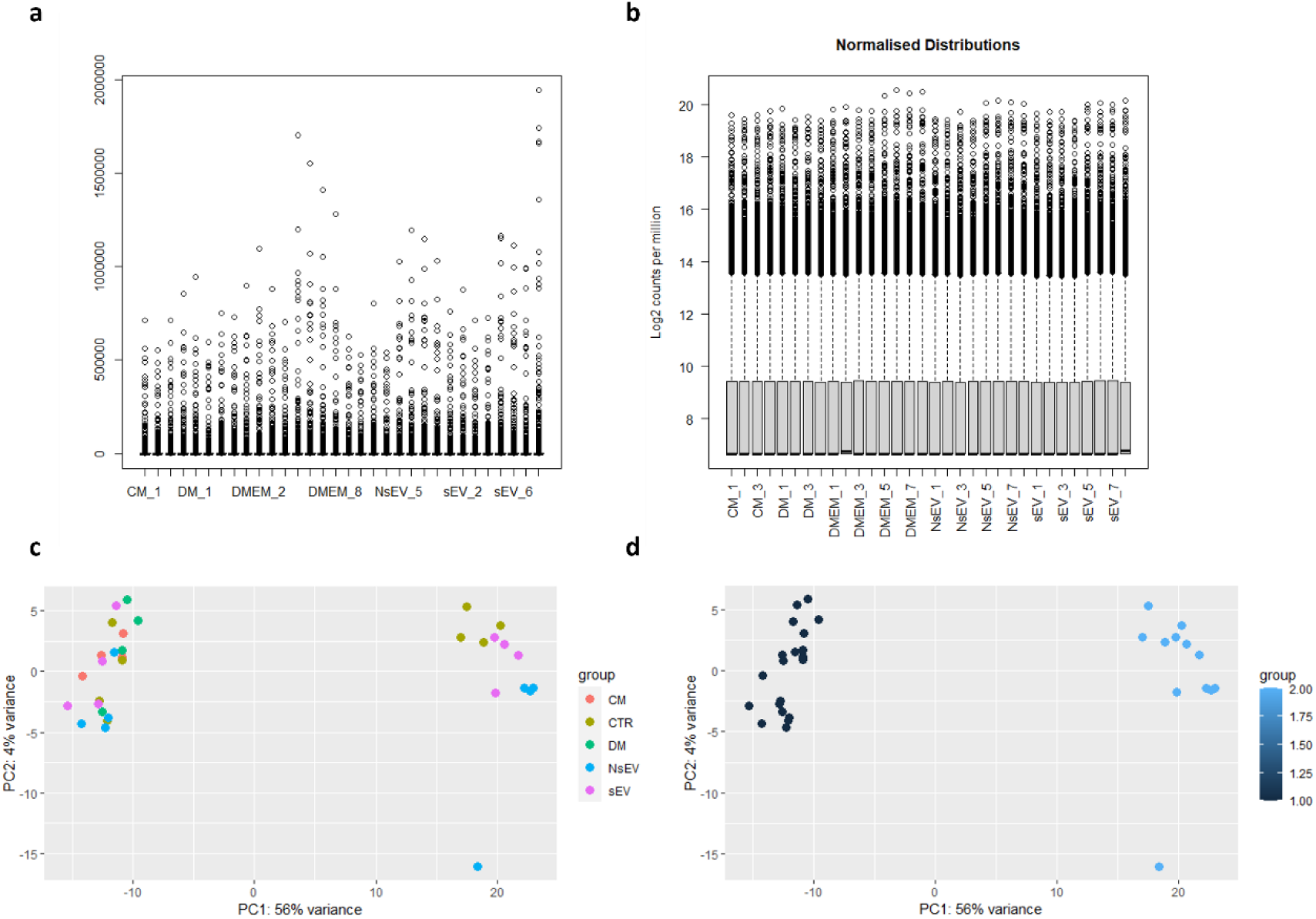
Quality control of reads and Principal Component Analysis (PCA). (a) Box plots displaying distribution of reads between samples. (b) Normalised distribution of reads. (c) PCA displaying major sources of variance between samples, according to their treatment. (d) PCA displaying major sources of variance between samples according to the batch, attesting for a batch effect.

**Supplementary Figure 5.**
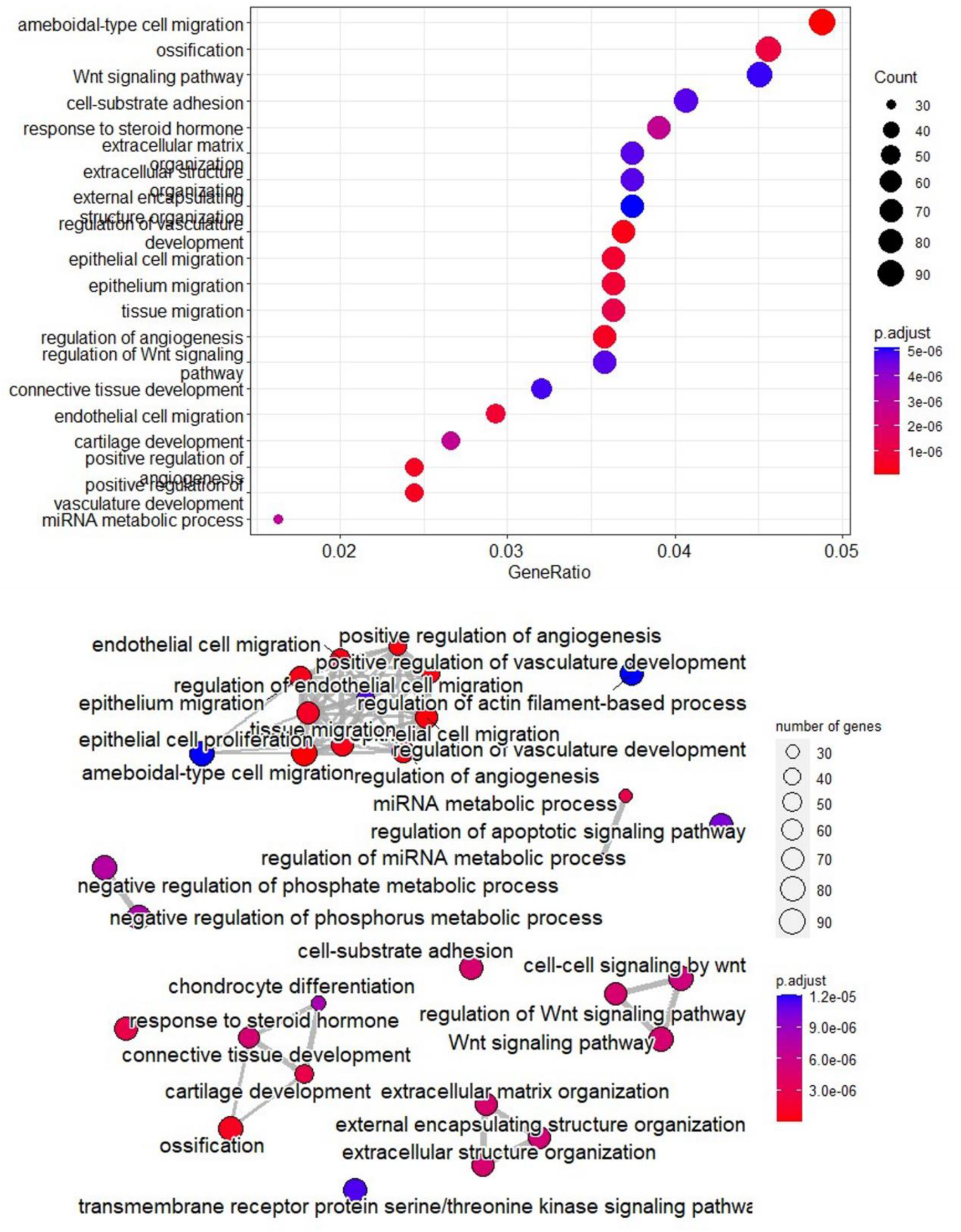
Gene Ontology Analysis and Enriched Pathways Clustering when comparing samples exposed to NsEVs and controls. (a) Pathways that have been significantly changed based on the number of genes within them that are differentially expressed. (b) Links between enriched pathways and how they cluster depending on common genes.

**Supplementary Table 1.** GLM ANOVA results for migration (cell covered area of the gap in μm2) of NHDFs in response to factors of migration time (Hour), three secretome fraction types from MSC (Treatment) and concentration of secretome (Concentration). DF is the abbreviation for Degree of Freedom.

| Variable | DF | F Value | P Value |
| --- | --- | --- | --- |
| Hour | 9 | 2032.78 | <0.001 |
| Treatment | 2 | 29.77 | <0.001 |
| Concentration | 2 | 11.48 | <0.001 |
| Hour*Treatment | 18 | 0.62 | 0.889 |
| Hour*Concentration | 18 | 2.10 | 0.005 |
| Treatment*Concentration | 4 | 11.38 | <0.001 |

**Supplementary Table 2.** Results of Tukey’s post-hoc multiple comparisons between different secretome fractions of MSC at different concentrations (e.g. MSC-sEV-1 means the group treated with MSC-sEV at 1 μg/mL) for their effects on migration behaviour of NHDFs. Mean Diff. is the abbreviation for Mean Difference.

| Tukey's multiple comparisons test | Mean 1 | Mean2 | Mean Diff. | Significance | P Value |
| --- | --- | --- | --- | --- | --- |
| Blank (DMEM) vs. MSC-sEV-1 | 672165 | 734542 | -62376 | **** | <0.0001 |
| Blank (DMEM) vs. MSC-sEV-30 | 672165 | 728205 | -56040 | **** | <0.0001 |
| Blank (DMEM) vs. MSC-CM-1 | 672165 | 668993 | 3172 | ns | >0.9999 |
| Blank (DMEM) vs. MSC-CM-30 | 672165 | 713417 | -41251 | *** | 0.0007 |
| Blank (DMEM) vs. MSC-NsEV-1 | 672165 | 665273 | 6893 | ns | 0.9927 |
| Blank (DMEM) vs. MSC-NsEV-30 | 672165 | 659940 | 12225 | ns | 0.8787 |
| MSC-sEV-1 vs. MSC-sEV-30 | 734542 | 728205 | 6336 | ns | 0.9954 |
| MSC-sEV-1 vs. MSC-CM-1 | 734542 | 668993 | 65548 | **** | <0.0001 |
| MSC-sEV-1 vs. MSC-CM-30 | 734542 | 713417 | 21125 | ns | 0.3304 |
| MSC-sEV-1 vs. MSC-NsEV-1 | 734542 | 665273 | 69269 | **** | <0.0001 |
| MSC-sEV-1 vs. MSC-NsEV-30 | 734542 | 659940 | 74602 | **** | <0.0001 |
| MSC-sEV-30 vs. MSC-CM-1 | 728205 | 668993 | 59212 | **** | <0.0001 |
| MSC-sEV-30 vs. MSC-CM-30 | 728205 | 713417 | 14789 | ns | 0.7458 |
| MSC-sEV-30 vs. MSC-NsEV-1 | 728205 | 665273 | 62933 | **** | <0.0001 |
| MSC-sEV-30 vs. MSC-NsEV-30 | 728205 | 659940 | 68265 | **** | <0.0001 |
| MSC-CM-1 vs. MSC-CM-30 | 668993 | 713417 | -44423 | *** | 0.0002 |
| MSC-CM-1 vs. MSC-NsEV-1 | 668993 | 665273 | 3720 | ns | 0.9998 |
| MSC-CM-1 vs. MSC-NsEV-30 | 668993 | 659940 | 9053 | ns | 0.9698 |
| MSC-CM-30 vs. MSC-NsEV-1 | 713417 | 665273 | 48144 | **** | <0.0001 |
| MSC-CM-30 vs. MSC-NsEV-30 | 713417 | 659940 | 53477 | **** | <0.0001 |
| MSC-NsEV-1 vs. MSC-NsEV-30 | 665273 | 659940 | 5333 | ns | 0.9982 |

**Supplementary Table 3.** GLM ANOVA results for migration (cell covered area of the gap in μm2) of NHDFs in response to factors of migration time (Hour), two secretome fraction types from MSC (Treatment) and concentration of secretome (Concentration). DF is the abbreviation for Degree of Freedom.

| Variable | DF | F Value | P Value |
| --- | --- | --- | --- |
| Hour | 9 | 2143.94 | <0.001 |
| Treatment | 1 | 11.20 | 0.001 |
| Concentration | 4 | 31.06 | <0.001 |
| Hour*Treatment | 9 | 0.43 | 0.919 |
| Hour*Concentration | 36 | 2.20 | <0.001 |
| Treatment*Concentration | 4 | 4.75 | 0.001 |

**Supplementary Table 4.**
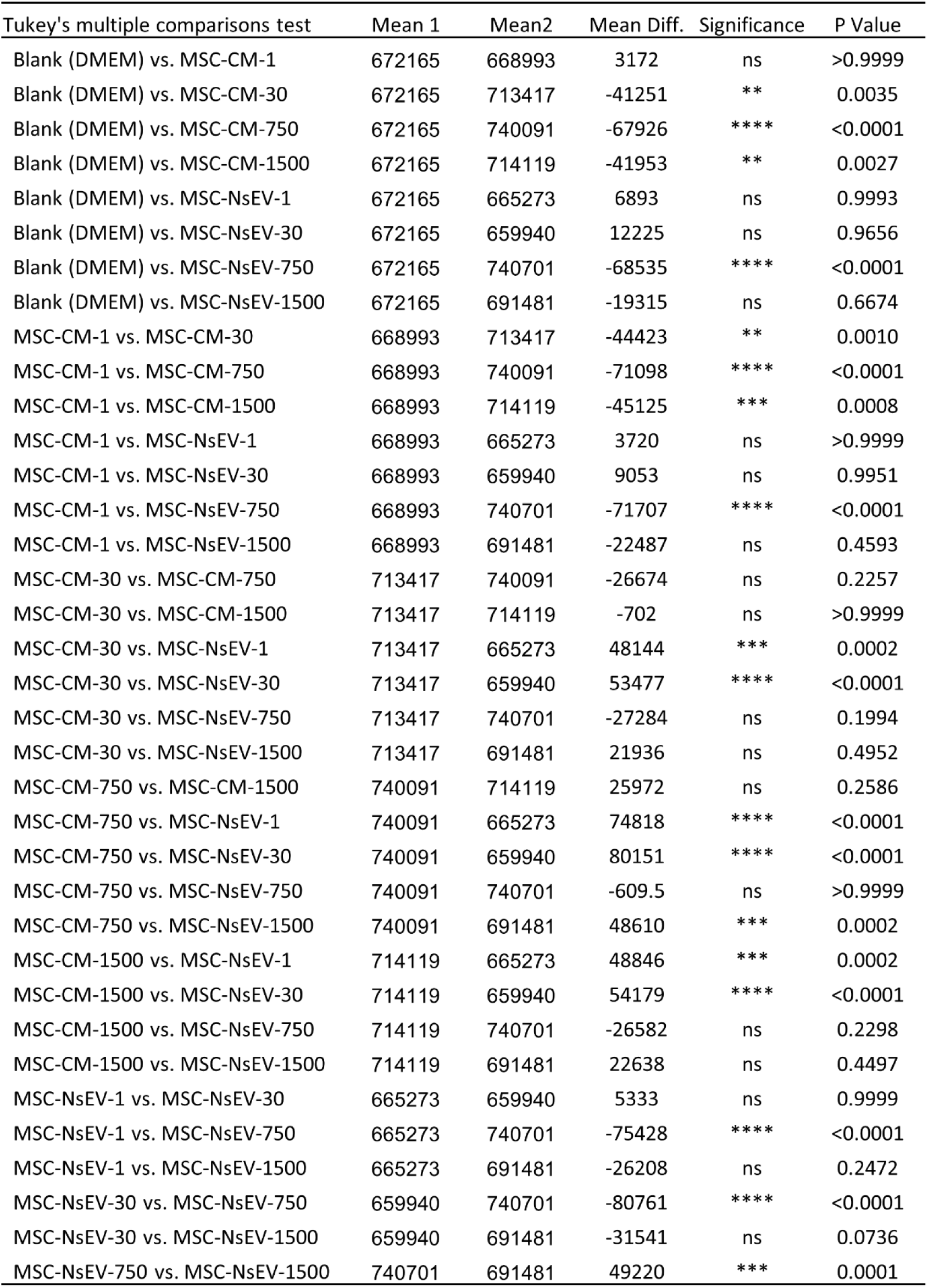
Results of Tukey’s post-hoc multiple comparisons between two secretome fractions of MSC at different concentrations (e.g. MSC-CM-1 means the group treated with MSC-CM at 1 μg/mL) for their effects on migration behaviour of NHDFs. Mean Diff. is the abbreviation for Mean Difference.

| Tukey's multiple comparisons test | Mean 1 | Mean2 | Mean Diff. | Significance | P Value |
| --- | --- | --- | --- | --- | --- |
| Blank (DMEM) vs. MSC-CM-1 | 672165 | 668993 | 3172 | ns | >0.9999 |
| Blank (DMEM) vs. MSC-CM-30 | 672165 | 713417 | -41251 | ** | 0.0035 |
| Blank (DMEM) vs. MSC-CM-750 | 672165 | 740091 | -67926 | **** | <0.0001 |
| Blank (DMEM) vs. MSC-CM-1500 | 672165 | 714119 | -41953 | ** | 0.0027 |
| Blank (DMEM) vs. MSC-NsEV-1 | 672165 | 665273 | 6893 | ns | 0.9993 |
| Blank (DMEM) vs. MSC-NsEV-30 | 672165 | 659940 | 12225 | ns | 0.9656 |
| Blank (DMEM) vs. MSC-NsEV-750 | 672165 | 740701 | -68535 | **** | <0.0001 |
| Blank (DMEM) vs. MSC-NsEV-1500 | 672165 | 691481 | -19315 | ns | 0.6674 |
| MSC-CM-1 vs. MSC-CM-30 | 668993 | 713417 | -44423 | ** | 0.0010 |
| MSC-CM-1 vs. MSC-CM-750 | 668993 | 740091 | -71098 | **** | <0.0001 |
| MSC-CM-1 vs. MSC-CM-1500 | 668993 | 714119 | -45125 | *** | 0.0008 |
| MSC-CM-1 vs. MSC-NsEV-1 | 668993 | 665273 | 3720 | ns | >0.9999 |
| MSC-CM-1 vs. MSC-NsEV-30 | 668993 | 659940 | 9053 | ns | 0.9951 |
| MSC-CM-1 vs. MSC-NsEV-750 | 668993 | 740701 | -71707 | **** | <0.0001 |
| MSC-CM-1 vs. MSC-NsEV-1500 | 668993 | 691481 | -22487 | ns | 0.4593 |
| MSC-CM-30 vs. MSC-CM-750 | 713417 | 740091 | -26674 | ns | 0.2257 |
| MSC-CM-30 vs. MSC-CM-1500 | 713417 | 714119 | -702 | ns | >0.9999 |
| MSC-CM-30 vs. MSC-NsEV-1 | 713417 | 665273 | 48144 | *** | 0.0002 |
| MSC-CM-30 vs. MSC-NsEV-30 | 713417 | 659940 | 53477 | **** | <0.0001 |
| MSC-CM-30 vs. MSC-NsEV-750 | 713417 | 740701 | -27284 | ns | 0.1994 |
| MSC-CM-30 vs. MSC-NsEV-1500 | 713417 | 691481 | 21936 | ns | 0.4952 |
| MSC-CM-750 vs. MSC-CM-1500 | 740091 | 714119 | 25972 | ns | 0.2586 |
| MSC-CM-750 vs. MSC-NsEV-1 | 740091 | 665273 | 74818 | **** | <0.0001 |
| MSC-CM-750 vs. MSC-NsEV-30 | 740091 | 659940 | 80151 | **** | <0.0001 |
| MSC-CM-750 vs. MSC-NsEV-750 | 740091 | 740701 | -609.5 | ns | >0.9999 |
| MSC-CM-750 vs. MSC-NsEV-1500 | 740091 | 691481 | 48610 | *** | 0.0002 |
| MSC-CM-1500 vs. MSC-NsEV-1 | 714119 | 665273 | 48846 | *** | 0.0002 |
| MSC-CM-1500 vs. MSC-NsEV-30 | 714119 | 659940 | 54179 | **** | <0.0001 |
| MSC-CM-1500 vs. MSC-NsEV-750 | 714119 | 740701 | -26582 | ns | 0.2298 |
| MSC-CM-1500 vs. MSC-NsEV-1500 | 714119 | 691481 | 22638 | ns | 0.4497 |
| MSC-NsEV-1 vs. MSC-NsEV-30 | 665273 | 659940 | 5333 | ns | 0.9999 |
| MSC-NsEV-1 vs. MSC-NsEV-750 | 665273 | 740701 | -75428 | **** | <0.0001 |
| MSC-NsEV-1 vs. MSC-NsEV-1500 | 665273 | 691481 | -26208 | ns | 0.2472 |
| MSC-NsEV-30 vs. MSC-NsEV-750 | 659940 | 740701 | -80761 | **** | <0.0001 |
| MSC-NsEV-30 vs. MSC-NsEV-1500 | 659940 | 691481 | -31541 | ns | 0.0736 |
| MSC-NsEV-750 vs. MSC-NsEV-1500 | 740701 | 691481 | 49220 | *** | 0.0001 |

**Supplementary Table 5.** GLM ANOVA results for migration (cell covered area of the gap in μm2) of NHDFs in response to factors of migration time (Hour), cell source of sEVs (sEV type) and concentration of sEVs (Concentration). DF is the abbreviation for Degree of Freedom.

| Variable | DF | F Value | P Value |
| --- | --- | --- | --- |
| Hour | 9 | 1844.09 | <0.001 |
| sEV type | 1 | 30.40 | <0.001 |
| Concentration | 4 | 9.28 | <0.001 |
| Hour*sEV type | 9 | 0.53 | 0.850 |
| Hour*Concentration | 36 | 1.70 | 0.007 |
| sEV type*Concentration | 4 | 2.49 | 0.042 |

**Supplementary Table 6.** Results of Tukey’s post-hoc multiple comparisons between different sEV types at different concentrations (e.g. MSC-sEV-0.01 means the group treated with MSC-sEV at 0.01 μg/mL) for their effects on migration behaviour of NHDFs. Mean Diff. is the abbreviation for Mean Difference.

| Tukey's multiple comparisons test | Mean 1 | Mean2 | Mean Diff. | Significance | P Value |
| --- | --- | --- | --- | --- | --- |
| Blank (DMEM) vs. MSC-sEV-0.01 | 672165 | 698860 | -26695 | ns | 0.3469 |
| Blank (DMEM) vs. MSC-sEV-0.1 | 672165 | 711355 | -39189 | * | 0.0225 |
| Blank (DMEM) vs. MSC-sEV-1 | 672165 | 734542 | -62376 | **** | <0.0001 |
| Blank (DMEM) vs. MSC-sEV-30 | 672165 | 728205 | -56040 | **** | <0.0001 |
| Blank (DMEM) vs. HDF-sEV-0.01 | 672165 | 672552 | -386.9 | ns | >0.9999 |
| Blank (DMEM) vs. HDF-sEV-0.1 | 672165 | 669409 | 2756 | ns | >0.9999 |
| Blank (DMEM) vs. HDF-sEV-1 | 672165 | 687378 | -15212 | ns | 0.9291 |
| Blank (DMEM) vs. HDF-sEV-30 | 672165 | 700293 | -28127 | ns | 0.2759 |
| MSC-sEV-0.01 vs. MSC-sEV-0.1 | 698860 | 711355 | -12494 | ns | 0.9778 |
| MSC-sEV-0.01 vs. MSC-sEV-1 | 698860 | 734542 | -35681 | ns | 0.0569 |
| MSC-sEV-0.01 vs. MSC-sEV-30 | 698860 | 728205 | -29345 | ns | 0.2230 |
| MSC-sEV-0.01 vs. HDF-sEV-0.01 | 698860 | 672552 | 26308 | ns | 0.3675 |
| MSC-sEV-0.01 vs. HDF-sEV-0.1 | 698860 | 669409 | 29451 | ns | 0.2187 |
| MSC-sEV-0.01 vs. HDF-sEV-1 | 698860 | 687378 | 11483 | ns | 0.9870 |
| MSC-sEV-0.01 vs. HDF-sEV-30 | 698860 | 700293 | -1432 | ns | >0.9999 |
| MSC-sEV-0.1 vs. MSC-sEV-1 | 711355 | 734542 | -23187 | ns | 0.5489 |
| MSC-sEV-0.1 vs. MSC-sEV-30 | 711355 | 728205 | -16851 | ns | 0.8785 |
| MSC-sEV-0.1 vs. HDF-sEV-0.01 | 711355 | 672552 | 38802 | * | 0.0251 |
| MSC-sEV-0.1 vs. HDF-sEV-0.1 | 711355 | 669409 | 41945 | * | 0.0100 |
| MSC-sEV-0.1 vs. HDF-sEV-1 | 711355 | 687378 | 23977 | ns | 0.5013 |
| MSC-sEV-0.1 vs. HDF-sEV-30 | 711355 | 700293 | 11062 | ns | 0.9898 |
| MSC-sEV-1 vs. MSC-sEV-30 | 734542 | 728205 | 6336 | ns | 0.9998 |
| MSC-sEV-1 vs. HDF-sEV-0.01 | 734542 | 672552 | 61989 | **** | <0.0001 |
| MSC-sEV-1 vs. HDF-sEV-0.1 | 734542 | 669409 | 65132 | **** | <0.0001 |
| MSC-sEV-1 vs. HDF-sEV-1 | 734542 | 687378 | 47164 | ** | 0.0018 |
| MSC-sEV-1 vs. HDF-sEV-30 | 734542 | 700293 | 34249 | ns | 0.0804 |
| MSC-sEV-30 vs. HDF-sEV-0.01 | 728205 | 672552 | 55653 | **** | <0.0001 |
| MSC-sEV-30 vs. HDF-sEV-0.1 | 728205 | 669409 | 58796 | **** | <0.0001 |
| MSC-sEV-30 vs. HDF-sEV-1 | 728205 | 687378 | 40828 | * | 0.0140 |
| MSC-sEV-30 vs. HDF-sEV-30 | 728205 | 700293 | 27913 | ns | 0.2860 |
| HDF-sEV-0.01 vs. HDF-sEV-0.1 | 672552 | 669409 | 3143 | ns | >0.9999 |
| HDF-sEV-0.01 vs. HDF-sEV-1 | 672552 | 687378 | -14826 | ns | 0.9386 |
| HDF-sEV-0.01 vs. HDF-sEV-30 | 672552 | 700293 | -27740 | ns | 0.2942 |
| HDF-sEV-0.1 vs. HDF-sEV-1 | 669409 | 687378 | -17969 | ns | 0.8341 |
| HDF-sEV-0.1 vs. HDF-sEV-30 | 669409 | 700293 | -30883 | ns | 0.1665 |
| HDF-sEV-1 vs. HDF-sEV-30 | 687378 | 700293 | -12915 | ns | 0.9727 |

**Supplementary Table 7.** GLM ANOVA results for proliferation (Folds of OD450 at Day 0) of NHDFs in response to factors of treatment time (Day), cell source of secretome (Cell Source) and secretome fraction type (Treatment) at concentration ≤ 30 μg/mL. DF is the abbreviation for Degree of Freedom.

| Variable | DF | F Value | P Value |
| --- | --- | --- | --- |
| Day | 2 | 407.25 | <0.001 |
| Cell source | 1 | 0.82 | 0.364 |
| Treatment | 2 | 3.10 | 0.046 |
| Day*Cell source | 2 | 0.23 | 0.796 |
| Day*Treatment | 4 | 1.29 | 0.273 |
| Cell source*Treatment | 2 | 0.34 | 0.709 |

**Supplementary Table 8.** Results of Tukey’s post-hoc multiple comparisons between different secretome fraction types at different concentrations (e.g. MSC-sEV-1 means the group treated with MSC-sEV at 1 μg/mL) for their effects on proliferation behaviour of NHDFs. Mean Diff. is the abbreviation for Mean Difference.

| Tukey's multiple comparisons test | Mean 1 | Mean2 | Mean Diff. | Significance | P Value |
| --- | --- | --- | --- | --- | --- |
| Blank (DMEM) vs. MSC-sEV-1 | 1.838 | 2.101 | -0.263 | ns | 0.9957 |
| Blank (DMEM) vs. MSC-sEV-30 | 1.838 | 2.461 | -0.624 | ns | 0.4988 |
| Blank (DMEM) vs. MSC-CM-1 | 1.838 | 2.486 | -0.648 | ns | 0.4419 |
| Blank (DMEM) vs. MSC-CM-30 | 1.838 | 2.754 | -0.917 | ns | 0.0750 |
| Blank (DMEM) vs. MSC-CM-750 | 1.838 | 4.547 | -2.710 | **** | <0.0001 |
| Blank (DMEM) vs. MSC-CM-1500 | 1.838 | 4.510 | -2.672 | **** | <0.0001 |
| Blank (DMEM) vs. MSC-NsEV-1 | 1.838 | 2.465 | -0.627 | ns | 0.4907 |
| Blank (DMEM) vs. MSC-NsEV-30 | 1.838 | 2.527 | -0.689 | ns | 0.3565 |
| Blank (DMEM) vs. MSC-NsEV-750 | 1.838 | 3.993 | -2.156 | **** | <0.0001 |
| Blank (DMEM) vs. MSC-NsEV-1500 | 1.838 | 4.142 | -2.304 | **** | <0.0001 |
| Blank (DMEM) vs. HDF-sEV-1 | 1.838 | 2.310 | -0.472 | ns | 0.8516 |
| Blank (DMEM) vs. HDF-sEV-30 | 1.838 | 2.145 | -0.307 | ns | 0.9942 |
| Blank (DMEM) vs. HDF-CM-1 | 1.838 | 2.455 | -0.617 | ns | 0.5144 |
| Blank (DMEM) vs. HDF-CM-30 | 1.838 | 2.771 | -0.933 | ns | 0.0655 |
| Blank (DMEM) vs. HDF-CM-750 | 1.838 | 3.901 | -2.063 | **** | <0.0001 |
| Blank (DMEM) vs. HDF-CM-1500 | 1.838 | 4.473 | -2.636 | **** | <0.0001 |
| Blank (DMEM) vs. HDF-NsEV-1 | 1.838 | 2.173 | -0.335 | ns | 0.9878 |
| Blank (DMEM) vs. HDF-NsEV-30 | 1.838 | 2.382 | -0.545 | ns | 0.6916 |
| Blank (DMEM) vs. HDF-NsEV-750 | 1.838 | 3.767 | -1.930 | **** | <0.0001 |
| Blank (DMEM) vs. HDF-NsEV-1500 | 1.838 | 4.302 | -2.464 | **** | <0.0001 |

## Acknowledgements

This work was supported by OSCAR’s research budget of Regenerative Medical Engineering Group and funding from Jiangsu Industrial Technology Research Institute (JITRI). The authors also would like to thank all members of both groups for their kind suggestions and comments, particularly Dr. Hui Wang for his contributions.

## Author contributions

Z.C. and H.Y. defined the project scope and provided research funding. H.Y, R.L and T.B.S. designed the experiments. R.L and T.B.S. performed the experiments, analysed data, and drafted the manuscript. Z.C. and H.Y. reviewed the manuscript. All authors approved the final version of the manuscript.

## Data availability statement

Data sets generated during this study are available from the corresponding authors upon reasonable request.

